# Equivariant neuronal populations enable simultaneous tuning and invariance

**DOI:** 10.1101/2024.08.02.606398

**Authors:** Judith Hoeller, Lin Zhong, Larissa Heinrich, Stephan Saalfeld, Marius Pachitariu, Sandro Romani

## Abstract

As we move through the world, we see the same visual scene from different perspectives. But how does the brain encode scene identity invariant to perspective, while remaining sensitive to these transformations? We propose a solution through equivariance, where perspective transformations induce structured changes in neuronal population responses. This framework implies a decomposition of population responses into orthogonal subspaces that are tuned and invariant. Testing our framework with large-scale neuronal recordings across four mouse visual cortical areas, we find that the equivariant structure is more pronounced in some higher-order areas (LM, AL) than in other areas (V1, RL). This equivariant structure accounts for the observed simultaneous increase in both population tuning and invariance. In comparison, early layers of an artificial neural network trained on image classification show similar structure, but later layers increase invariance at the cost of tuning. These results suggest equivariance is a principle to achieve flexible computations with neuronal populations.

---

A fundamental challenge in vision neuroscience is understanding how the brain constructs stable percepts from retinal images that are constantly being transformed by our own movements. Despite this perpetual flux, our perception of scene and object identity remains remarkably stable, even as we simultaneously detect the transformations themselves. This raises a central question: How does the visual system achieve the dual goal of transformation-independent (“invariant”) identification and transformation-dependent (“tuned”) detection (**Fig 1a**)?

**Fig 1.**
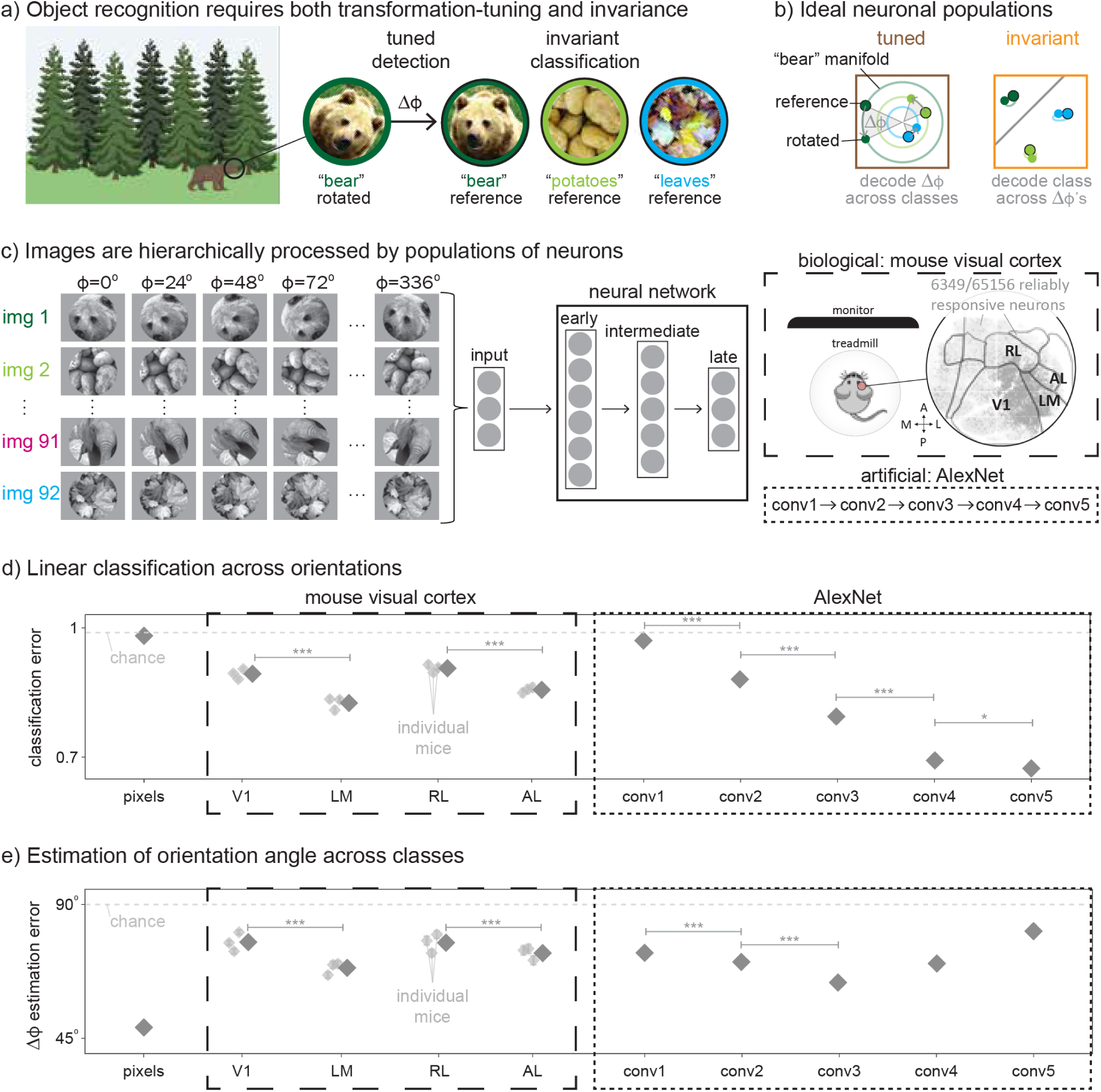
Population decoding of image identity and relative orientation across mouse visual cortical areas and AlexNet layers. **a)** Schematic of object recognition, which requires detection, i.e., rotating an image by an angle Δϕ to align with a reference image, and classification. The left-most image was adapted from “Tree (coniferous, forest)” and “Brown Bear (Ursus arctos)”, by BioRender.com (2024). **b)** Ideal neuronal responses that enable a simple decoding of Δϕ across class (left) or class across Δϕ (right); colored by class as in a. **c)** A feedforward neural network processing 92 images, each rotated 15 times by Δϕ = 24°. The upper-right panel includes a schematic of the experimental setup. In the circular inset, light gray dots indicate the positions of imaged neurons, darker gray dots the reliably responsive neurons, and black lines estimated area boundaries. **d)** Classification error of a linear SVM, which decodes class across Δϕ from neuronal responses. Shown is the classification error for held-out data (mean ± sem across 1380 samples). **e)** Estimation of Δϕ from neuronal responses using a simple relative orientation decoder. Shown is the absolute estimation error in Δϕ for held-out data (mean ± sem across 1288 samples). In d-e, sample sizes in mouse visual cortical areas are 3 times larger (for 3 mice; results for individual mice are shown as smaller diamonds), results are averaged over at most 100 independent random samples of N = 250 neurons, error bars are smaller than dot sizes, and stars reveal statistically significant decreases.

The traditional view is that this is achieved at the level of single neurons, where neurons in early areas are tuned and those in later areas are invariant. But pioneering studies^1–4^ have shown that neuronal populations in primate visual cortex are instead jointly tuned and invariant, with earlier areas exhibiting stronger tuning and later areas showing increased invariance, yet without fully losing sensitivity to image transformations. Notably, object identity as well as transformation variables could be decoded from the same neuronal population, and decoding of both improved across the visual hierarchy. Yet no existing framework specifies what geometric structure neuronal population responses should have to simultaneously support invariant classification and tuned detection.^5,6^

Two prominent hypotheses address how invariance might emerge in a population code. The untangling hypothesis^1–3^ proposes a gradual reformatting of object manifolds—the collection of population responses to an object and its transformations—to increase linear separability across the visual hierarchy. In contrast, the equivariance hypothesis^7,8^ posits that image transformations lead to structured, linear transformations of population responses. Since geometric transformations such as in-plane rotations act as linear transformations on pixels, equivariance is one of the simplest and most natural ways for population responses to change. Invariance arises as a special case of equivariance, which may be achieved through pooling.^9^ Whereas untangling focuses on the linear separability of object identity from population responses, equivariance emphasizes the geometric structure within those responses.

To investigate tuning and invariance empirically, we recorded neuronal responses in mice, which offer large-scale neuronal access^10,11^ and exhibit complex visual behaviors relevant for object recognition.^12–15^ We focused on in-plane rotations to probe the population code in a controlled manner (**Fig 1c**). Rotations have nontrivial geometric consequences for population responses: because a 360° rotation returns the original image, object manifolds form closed loops. Consistent with primate studies,^1–4^ overlapping neuronal populations supported the simple decoding of image identity and rotation angle (relative orientation), with decoding performance improving from V1 to higher-order areas (LM and AL but not RL).

To interpret these findings, we developed a geometric framework in which equivariance and untangling jointly support simple decoding of both image identity and relative orientation. We show that equivariance induces a decomposition of population responses into orthogonal subspaces: tuned subspaces in which object manifolds form closed loops (**Fig 1b** left), and invariant subspaces in which manifolds collapse to points (**Fig 1b** right). This organization provides a principled way for tuning and invariance to coexist within a single population code, making it simple for downstream neurons to separately read-out image identity and relative orientation.

We further applied this geometric framework to an artificial visual system solely trained on object classification. In AlexNet^16,17^ trained on ImageNet,^18^ early-to-intermediate layers showed equivariant structure resembling that of mouse visual cortex. However, in later layers, equivariance became increasingly biased toward invariance, and relative orientation decoding degraded. Thus, training for object classification alone does not guarantee the equivariant structure required for detection.

Finally, equivariance has received growing attention in artificial neural networks, where architectural constraints can guarantee invariances by design.^19–22^ Convolutional neural networks are among the best predictive models of biological visual responses,^23–28^ and incorporating equivariance-inspired constraints can further improve prediction.^29,30^ Our approach provides a general framework for characterizing equivariant structure of population responses in biological and artificial neural networks. Together, our findings synthesize untangling and equivariance into a unified framework of how structured neuronal populations can support invariant identification and tuned detection.

## RESULTS

### Population decoding of image identity and relative orientation in mouse visual cortex

To investigate how object identity and relative orientation are represented across mouse visual cortical areas, we performed large-scale two-photon calcium imaging on 3 awake mice head-fixed on a spherical treadmill (**Fig 1c** top-right). Mice passively viewed naturalistic, grayscale images at multiple orientations, interleaved with a gray screen—an experimental paradigm originally developed in primates^1,4,31,32^ and adapted for rodents,^14,33–36^ to probe fast visual processing of as many stimuli as possible. Mice have been shown to behaviorally discriminate complex visual stimuli,^14,15,37^ including naturalistic scenes and textures similar to ours, validating that such stimuli engage the mouse visual system in a meaningful way.

The stimulus set consisted of 92 image identities (46 scenes and 46 textures) presented across 15 evenly spaced orientations (**Fig 1c** left), allowing us to separately examine representations of identity and relative orientation. This design leverages the observation that much of the difficulty of object recognition arises from variability due to transformations rather than from differences between identities within the same object class.^2,9^ Each of 90 image identities was displayed twice (img 1-90), and the other 2 identities eighty times (img 91-92), which allowed us to maximize the number of stimulus presentations while maintaining 2 identities with a more accurate estimate of the stimulus-driven neuronal response. Images were randomly displayed on either the front or left monitor. In the main text, we focus our analyses on the front monitor; results for the left monitor are presented in **Supplementary Fig 5**.

For each mouse, we recorded more than 50,000 neurons across four retinotopic visual cortical areas (retinotopies in **Extended Data Fig 1a**), including V1 (primary), LM (lateromedial), RL (rostrolateral) and AL (anterolateral). It is known that V1, LM, and AL are required for orientation discrimination (RL was not tested).^38^ These areas lie in both the putative ventral (LM) and dorsal (RL, AL) streams and are at different stages of the visual hierarchy (V1 comes before LM and RL which are before AL).^36,39–41^ We note that the hierarchical organization of visual cortex is shallower and less established in mice than in primates.^40,42^

Reliably responsive neurons were mainly found in one part of visual cortex (**Fig 1c** top-right), consistent with the retinotopic location of the front screen. The fraction of such neurons over the total number in each area is 0.16 ± 0.02 in V1, 0.28 ± 0.01 in LM, 0.07 ± 0.01 in RL, and 0.14 ± 0.01 in AL (mean ± standard error of the mean (sem) across 3 mice). Other single-neuron response properties, such as trial-to-trial reliability, orientation selectivity and discrimination indices, were similar across areas (**Extended Data Fig 1b**). At both the single-neuron and population level, responses varied with image identity and relative orientation in complex ways (**Supplementary Fig 1-2**).

To test untangling, we probed to what degree image identity could be linearly decoded from neuronal responses in each visual area, in a way that generalizes across orientations. Similar to previous works,^4,43^ we trained a linear support vector machine (SVM)—a high-dimensional version of the linear decoder illustrated in **Fig 1b** right—to classify images across multiple orientations. The performance of the SVM is given by the classification error, *Err*, on a single held-out orientation. With 92 image classes, chance performance corresponds to *Err* ≈ 0.99.

To quantify how object identity is represented across visual areas, we randomly sampled a fixed number of neurons from each area (*N* = 250, limited by AL), following established practice in population decoding studies.^1–3,33,44,45^ As a baseline, decoding on pixels was near chance-level (*Err* = 0.9816 ± 0.0007 is the mean±sem across 15 individually held-out orientations and 92 classes), confirming that image identity cannot simply be recovered from the pixel representation. In contrast, decoding from neuronal population responses was consistently above chance in all visual areas (**Fig 1d**), with *Err* = 0.893 ± 0.002 in V1, 0.825 ± 0.004 in LM, 0.906 ± 0.003 in RL, and 0.856 ± 0.005 in AL (pooled across 3 mice), and was similar across scenes and textures as well as the held-out orientation (**Extended Data Fig 2a-b**).

Although decoding accuracy was modest with *N* = 250 neurons, these values are comparable to those reported in primate visual cortex under similar experimental conditions (Ref. [4]). Consistent with this work in primate visual cortex, classification performance changed systematically across mouse visual cortical areas, indicating structured differences in how image identity is represented. A direct geometric correlate of improved classification is that the distance between mean neuronal responses to held-out image identities—normalized by within-identity variability—increased systematically from V1 to higher-order areas (**Extended Data Fig 5b**). Errors decreased with *N* while the area ranking is preserved (**Extended Data Fig 3b**), indicating that performance at *N* = 250 reflects subsampling rather than absence of representational structure. At maximum *N*, V1 and LM had similar errors, suggesting LM contains comparable information with fewer neurons.

We next tested to what degree relative orientation could be decoded from the same neuronal population responses, in a way that generalizes across identities. Neurons in mouse visual cortical areas, especially in V1 but also in higher-order areas, are known for their orientation-tuning.^46,47^ To assess the decodability of relative orientation at the level of neuronal populations, we developed a simple decoder that takes population responses to an image at two orientations, ϕ_1_and ϕ_2_, and estimates the relative orientation, Δϕ = ϕ_2_ − ϕ_1_. We denote the output of this simple relative orientation decoder by Δϕ̂.

The idea of the relative orientation decoder comes from the observation that if population responses lie on concentric circles in a shared plane (**Fig 1b** left), then Δϕ would be simply decodable across all identities. We confirmed that, for each image identity, population responses were approximately equidistant from their mean (**Extended Data Fig 5a**), consistent with this geometric structure. We defined the shared plane by the first two principal components (PCs) of mean-subtracted population responses to 91 classes. We tested the relative orientation decoder on population responses to the single held-out class at ϕ_1_ = 0° and ϕ_2_ = 24°, …, 336°, and evaluated its performance using the estimation error, |Δϕ̂ − Δϕ|. The shared plane defined by PCs may not be ideal, hence the obtained |Δϕ̂ − Δϕ| should be viewed as an upper bound, and chance performance is |Δϕ̂ − Δϕ| ≈ 90°. We note that the observed relative orientation decoder is independent of the reference orientation ϕ_1_ (**Supplementary Fig 3**).

Relative orientation decoding performance showed systematic differences across areas, mirroring the trends observed for identity decoding. The estimation error was lower in LM and AL than in V1 and RL (V1: |Δϕ̂ − Δϕ| = 77.3° ± 0.4°, LM: 68.6° ± 0.6°, RL: 77.1° ± 0.7°, AL: 73.6° ± 0.8° across the 14 held-out orientations and 92 classes, pooled across 3 mice; **Fig 1e**). The decoding error for relative orientation was lower for scenes than for textures; smaller angles were also estimated more accurately than larger ones (**Extended Data Fig 2a,c**). As expected, the relative orientation decoder performed better on pixels (48.6° ± 0.7°), reflecting the fact that image rotations correspond to exact geometric transformations in pixel space. Pixel performance was largely limited by subsampling (250 of 50,176 pixels).

To test whether identity and orientation information were carried by segregated neuronal subpopulations, we compared the decoder weights (**Extended Data Fig 6**). The neurons contributing most to identity decoding substantially overlapped with those contributing to relative orientation decoding, consistent with theoretical arguments that mixed selectivity in high-dimensional populations facilitates flexible readout.^48–50^

These results demonstrate that neuronal populations in each mouse visual cortical area support above-chance decoding of both image identity and relative orientation, changing systematically across areas. Decoding performances at *N* = 250 were strongly limited by subsampling; when all reliably responsive neurons were used, *Err* = 0.659 ± 0.007 and |Δϕ̂ − Δϕ| = 67.8° ± 0.8° in V1 (**Extended Data Fig 4a**).

### Population decoding of image identity and relative orientation in AlexNet

Having established simple, simultaneous decoding in mouse visual cortex, we next examined whether similar decoding patterns emerge in an artificial neural network trained solely on image classification. We analyzed AlexNet,^16,17^ a deep, convolutional neural network, trained on ImageNet.^18^ While the organization of mouse visual cortex differs substantially from that of deep convolutional networks and does not form a strict feedforward hierarchy, AlexNet provides a controlled visual system in which changes in the population code across successive processing stages can be examined.

AlexNet comprises five convolutional layers (conv1-5) with increasing receptive field sizes. We adjusted the input image size such that receptive field sizes in conv1 were comparable to those in mouse V1. We focused our analyses on the convolutional layers, which preserve spatial structure and thus provide a closer functional analogue to visual cortical areas.^24,27^ To simulate trial-to-trial variability, we added independent Gaussian noise to each artificial neuronal response (**Online Methods**).

When we decoded image identity from *N* = 250 artificial neuronal responses in each convolutional layer, classification performance improved across layers (conv1: *Err* = 0.970 ± 0.001, conv2: 0.885 ± 0.004, conv3: 0.789 ± 0.007, conv4: 0.697 ± 0.009, conv5: 0.70 ± 0.01; **Fig 1d**). Relative orientation decoding improved up until intermediate layers (conv1: |Δϕ̂ − Δϕ| = 73.7° ± 0.5°, conv2: 70.6° ± 0.7°, conv3: 65.2° ± 0.9°), but then worsened in late layers (conv4: 75° ± 1°, conv5: 81° ± 1°; **Fig 1e**).

Decoding performances for both image identity and relative orientation in mouse V1 and RL fall around AlexNet conv1-2, and those in LM and AL around conv2-3 (Fig 1d-e). The divergence between mouse visual cortex and AlexNet appears in conv4–5, where invariance is prioritized at the expense of relative orientation tuning. Using a multiplicative gain model fit to our data produced similar decoding trends, except that relative orientation estimation progressively degraded across layers (**Extended Data Fig 7d-f**). We also trained AlexNet variants on different objectives (**Extended Data Fig 8**). The networks that did not include classification—orientation-only and randomly initialized—showed little improvement in classification performance across layers and did not resemble mouse visual cortex. The three variants trained on classification (default, rotation-augmented, and multi-task) all showed strong improvements across layers, resembling mouse visual cortex in conv1-3. Classification training therefore appears to be part of what produces cortex-like structure in early-to-intermediate AlexNet layers.

### Rotation-equivariant populations support both tuning and invariance

What is the structure of population codes that support simple decoding of both image identity and relative orientation, as observed in mouse LM, AL and AlexNet conv3? To address this, we analyzed the geometry of object manifolds under planar rotations using tools from mathematical group theory. While object manifolds must form loops under image rotations, their precise shapes and mutual relationships could be unstructured (**Fig 2a**). Here, we show that equivariance induces a decomposition of population responses into orthogonal subspaces that separately support invariant classification and tuned detection. To keep the discussion accessible, most mathematical details are deferred to the **Supplementary Information**.

**Fig 2.**
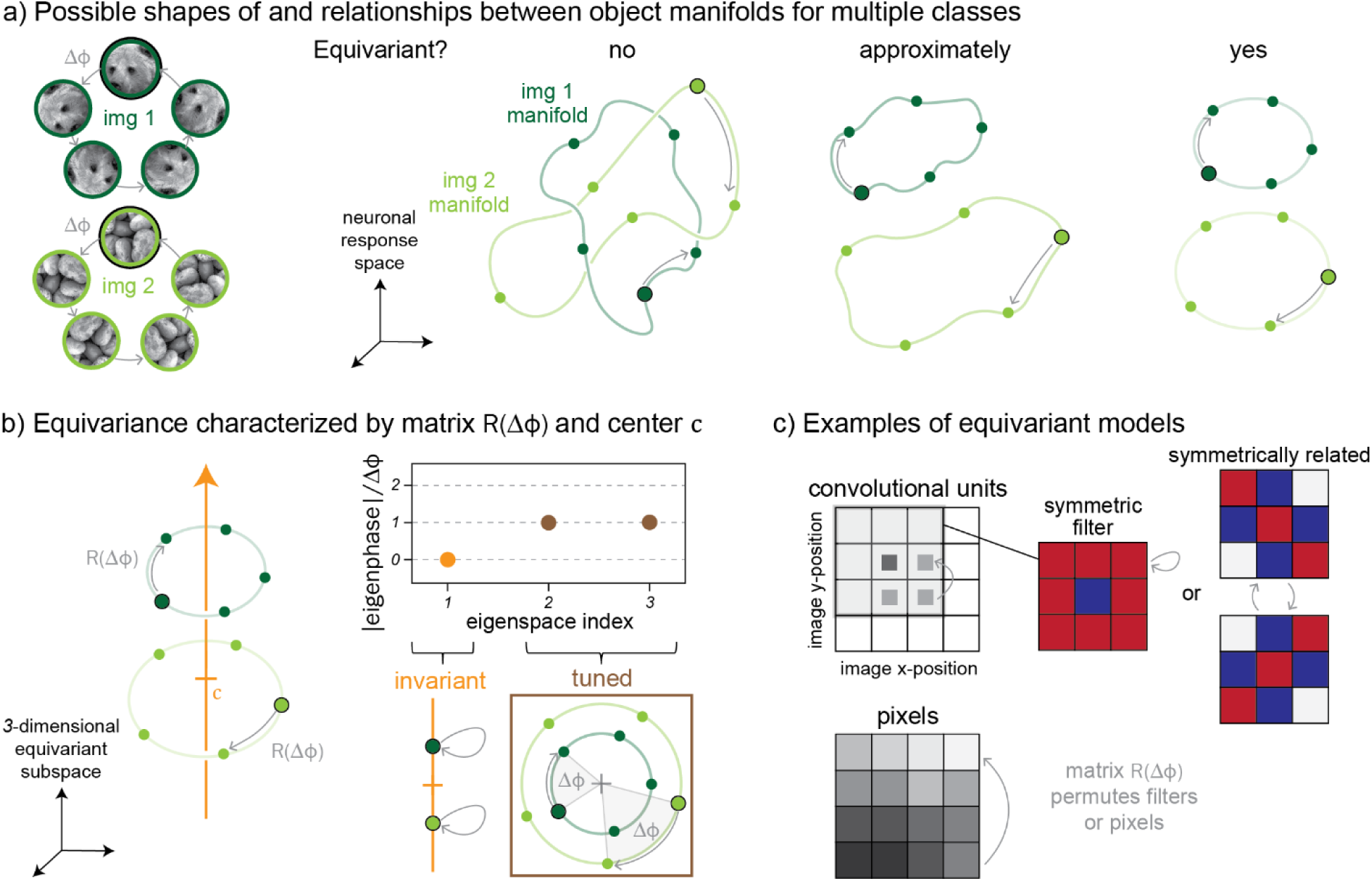
Rotation-equivariant population responses decompose into orthogonal invariant and tuned subspaces. **a)** Image rotations cause neuronal population responses to trace out closed loops (object manifolds), one per image. In general, these loops can have any shape (“no”). When responses are rotation-equivariant, the loops become circles aligned along shared axes (“yes”). When responses are approximately equivariant, they deviate from this strongly constrained structure (“approximately”). **b)** As the image rotates by an angle Δϕ, population responses rotate about a center c. The rotation axis (orange) is unchanged by rotation—the invariant direction—while the other two (brown) form a plane in which responses rotate by the same angle Δϕ—the tuned directions. **c)** Two examples of fully known equivariant models. Top: a linear receptive-field model with convolutional units on a 2×2 grid of image positions; for the darker square, the 3×3 filter is shown on the image (blue: negative weights, red: positive, white: zero). Bottom: the pixel model, where “responses” are grayscale pixel values. In both cases, an image rotation acts as a permutation of the units.

Rotation-equivariant populations exhibit a special subspace structure: they contain (1) invariant subspaces, in which object manifolds collapse to points, and (2) tuned subspaces, where manifolds form concentric circles (**Fig 2b**). These subspaces are orthogonal. The invariant subspace is ideal for supporting linear decoding of image identity across orientations, while the tuned subspace is ideal for simple decoding of relative orientation across identities. In general, each invariant direction can be thought of as a rotation axis, and each pair of tuned directions as a plane in which population responses rotate by the same angle as the image. In the combined subspace, formed by the invariant and tuned directions, object manifolds form concentric circles that are displaced along the invariant directions.

The subspace structure is guaranteed to exist in a particular basis of equivariant population responses (see **Supplementary Information**). In that basis, rotation-equivariance means that population responses to an image and to its rotated version are related by an orthogonal transformation *R*(Δϕ) (Fig 2b left), independent of image identity. Constraints from the group structure of planar rotations imply that the eigenvalues of *R*(Δϕ) have the form *e*^*im*Δϕ^ where *m* is an integer; we refer to *m*Δϕ as the eigenphase and order eigenspaces by increasing absolute value of *m*. Eigenspaces with *m* = 0 form the “invariant” subspace, where responses do not change with rotations, and eigenspaces with *m* = ±1 form the “tuned” subspace, where responses rotate by Δϕ in a plane (**Fig 2b**). Their dimensions are denoted by *d*_*i*_ and *d*_*t*_, respectively, with *d*_*t*_ always even. Higher-order eigenspaces corresponding to larger *m* may exist, analogous to higher-order Fourier modes, but are not relevant for the simple decoders used in this work.

Equivariance can take various forms. In the fully invariant case, *d*_*i*_ = *d* and *d*_*t*_ = 0, *R*(Δϕ) is the identity matrix and object manifolds are just points. In the fully tuned case, *d*_*i*_ = 0 and *d*_*t*_ = *d*, *R*(Δϕ) acts like 2D rotations by Δϕ and object manifolds are just circles. The pixel representation (**Fig 2c** bottom) provides an example where both invariant and tuned subspaces are present. As *R*(Δϕ) permutes pixels, pixels that lie on a fixed distance from the center are invariant for certain weighted combinations and are tuned for others (**Extended Data Fig 9**). Even though the invariant subspace exists in this case, it encodes low-level image statistics rather than image identity. Linear receptive-field models with symmetric or symmetrically related kernels exhibit similar equivariant structure (**Fig 2c** top; also Ref. [9]).

Relative orientation can be decoded directly from the tuned subspace. Within each pair of tuned directions and across all image identities, the action of *R*(Δϕ) reduces to a planar rotation by Δϕ. Consequently, the angular distance between two points on the projected object manifold directly reflects their difference in image orientation (**Fig 2b** bottom-right). As a result, the relative orientation decoder introduced earlier should perform perfectly within any pair of tuned directions, provided that *d*_*t*_ > 0.

On the other hand, image identity is simpler to decode in the invariant subspace than in the full space (see also ref. [51]), because variability due to orientation has been factored out. However, object manifolds projected to the invariant subspace do not necessarily encode identity, as we have seen for pixels. Linear separability depends on how points are distributed within the invariant subspace. For randomly distributed points, increasing the dimensionality *d*_*i*_ generically improves linear separability.^52^ For non-random distributions, points can be linearly separable even in low dimensions. (For example, points that are uniformly distributed on a circle are linearly separable.) Thus, an additional mechanism such as untangling within the invariant subspace is crucial for improving classification performance.

Equivariance provides a geometric framework for understanding how population responses can simultaneously support invariant classification (together with untangling) and tuned detection. Classification performance is determined by the dimensionality of the invariant subspace and by how neuronal responses within it are arranged (untangling), whereas the simple relative orientation decoder should perform perfectly within the tuned subspace if it exists.

### A computational method to quantify rotation-equivariance in population responses

To estimate rotation-equivariance from data, we developed a computational method that finds an equivariant subspace in population responses, along with a rotation matrix *R*(Δϕ) and a rotation center *c*. At its core, the method involves three simple, linear-algebraic steps: (1) projection of approximately whitened population responses to a low-dimensional subspace, (2) estimation of a single *R*(Δϕ) and *c* that best map responses before and after image rotation, and (3) eigendecomposition of *R*(Δϕ) which identifies the invariant and tuned subspace. This method does not rely on any particular network architecture or single-neuron tuning properties. The method takes as input *T* population response pairs, each obtained from two consecutive image rotations of the same image identity, sampled across multiple identities (**Fig 3a**; *T* = 1,380).

**Fig 3.**
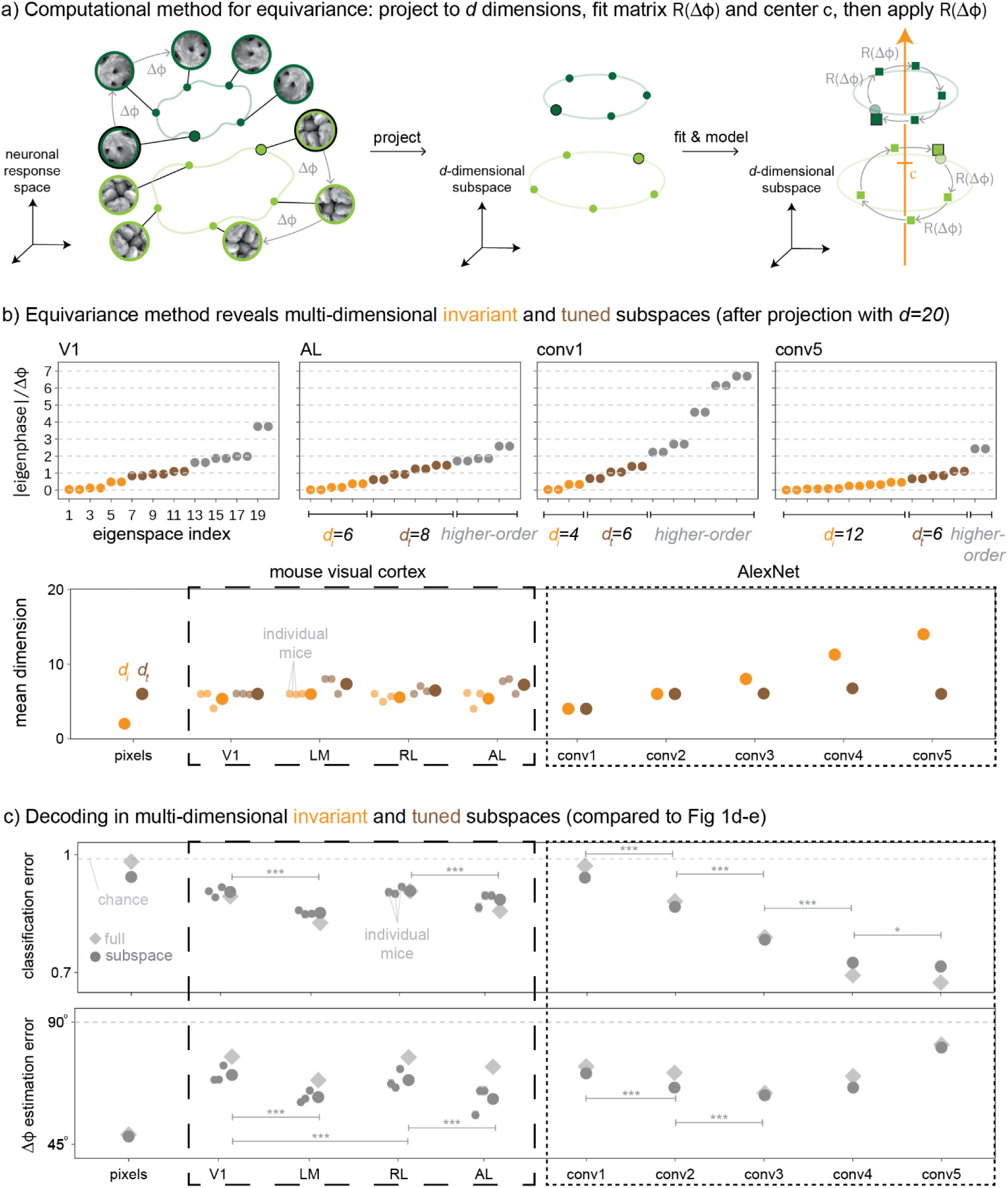
Computational method recovers equivariant subspace structure and recapitulates decoding trends across mouse visual areas and AlexNet layers. **a)** Our computational method consists of a projection to a *d*-dimensional subspace and estimation of a *d* × *d* rotation matrix *R*(Δϕ) and *d*-dimensional center *c*. Starting with neuronal responses to the reference image for each class (black border), we model responses to all rotated images by subsequent applications of *R*(Δϕ) (squares on the right; squares with black borders indicate predictions after rotation by Δϕ = 360°). **b)** Top: Analogue of Fig 2b right, with *R*(Δϕ) estimated in example areas (trained on img 1-90, 250 random neurons in each area, V1 and AL from mouse 2). Integer multiples of Δϕ are shown as gray dashed lines and dots are gray if the closest integer multiple is larger than 1. Bottom: Summary of invariant and tuned subspace dimensions, *d*_*i*_ and *d*_*t*_, respectively. **c)** Analogue of Fig 1d-e (diamonds) but for projections to invariant (top) and tuned (bottom) subspaces (dots). In b bottom and c, sample sizes in mouse areas are 3 times larger (results for individual mice are shown as smaller dots), results are averaged over at most 100 independent random samples of N = 250 neurons, error bars are smaller than dot sizes, and stars reveal statistically significant decreases across areas.

If *T* is smaller than the number of neurons (e.g. all neurons in V1), the empirical estimate of *R*(Δϕ) is not full-rank and thus cannot be orthogonal. To address this, we first projected whitened population responses into a *d*-dimensional subspace with *d* ≤ *T*. Specifically, the *d*-dimensional subspace was defined based on maximizing covariances between response pairs, like in canonical correlation analysis.

Within the *d*-dimensional subspace, we estimated a single *d* × *d* rotation matrix *R*(Δϕ) using a variant of orthogonal Procrustes analysis.^53^ If image rotations are uniformly sampled over 360°, as in our case, the rotation center *c* is simply equal to the mean projected population response.

Given the *d*-dimensional subspace projection, the rotation matrix *R*(Δϕ), and the rotation center *c*, we defined an equivariance model that predicts the entire object manifold from the population response to a single reference orientation (**Fig 3a** right). Object manifolds predicted by the equivariance model are highly constrained: they are points if *d*_*i*_ = *d*, concentric circles if *d*_*t*_ = *d*, and combinations thereof, i.e., precisely aligned circles, if *d*_*i*_ and *d*_*t*_ > 0.

Previous approaches to estimate equivariance in neuronal recordings were indirect, i.e., by fitting the data to an equivariant or equivariance-promoting artificial neural network.^29,30^ To our knowledge, there exist few computational methods that quantify equivariance and they only apply to artificial neural networks.^54,55^ In contrast, our method is agnostic to the underlying network architecture (biological or artificial). We validated our method on pixels, where the equivariant structure is fully known (as discussed before), and found that it accurately recovers the expected invariant and tuned subspaces (**Extended Data Fig 4b**; *N* = 4000). The AlexNet variants further confirm our method detects meaningful differences in equivariant structure across networks with different training objectives (**Extended Data Fig 8**).

### Structure of rotation-equivariance in mouse visual cortex and AlexNet

To identify the structure of rotation-equivariance in mouse visual cortical areas and AlexNet layers, we applied our computational method to a fixed number of neurons in each area and layer (*N* = 250, as before). We chose the subspace dimension *d* = 20, as model performances saturated around this dimension in all cases (**Extended Data Fig 10**).

We estimated the dimension of the invariant and tuned subspaces, *d*_*i*_ and *d*_*t*_, by counting discretized eigenphases Δθ_*j*_ of the estimated rotation matrices *R*(Δϕ) (**Fig 3b**). In mouse visual cortex, *d*_*i*_ remained roughly constant across areas (between *d*_*i*_ = 5.37 ± 0.05 and 5.97 ± 0.03 is the mean ± sem across 46 choices of 90 training classes, pooled across 3 mice), while *d*_*i*_ strongly increased across AlexNet layers, from *d*_*i*_ = 4.0 ± 0.0 in conv1 to *d*_*i*_ = 13.6 ± 0.1 in conv5. On the other hand, *d*_*t*_ was nonzero in all areas and layers—ranging between 5.79 ± 0.09 and 7.33 ± 0.03 in mouse visual cortex, and between 5.4 ± 0.1 and 7.1 ± 0.1 in AlexNet.

For equivariant neuronal responses, classification performance depends on *d*_*i*_ and the projection of neuronal responses to the invariant subspace while relative orientation decoding is determined by the presence of the tuned subspace. Indeed, decoding in those subspaces (**Fig 3c**) shows the same decoding trends as before (**Fig 1d-e**), with the relative orientation decoder performing consistently better in mouse visual cortex (AUC values between 0.53 and 0.60, see **Online Methods** for more effect size measures). This is because our computational method identifies the ideal shared planes for decoding relative orientation (pairs of tuned subspaces) while PCA does not.

To visualize the high-dimensional object manifolds and assess their geometric structure, we projected held-out neuronal responses to the tuned and invariant subspaces (**Fig 4a**). We chose the pair of tuned and invariant dimensions where the eigenphases were closest to integer multiples of Δϕ. These projections are more interpretable than those using PCs (**Supplementary Fig 1**), which generically mix neuronal variance due to image identity and variance due to orientation.

**Fig 4.**
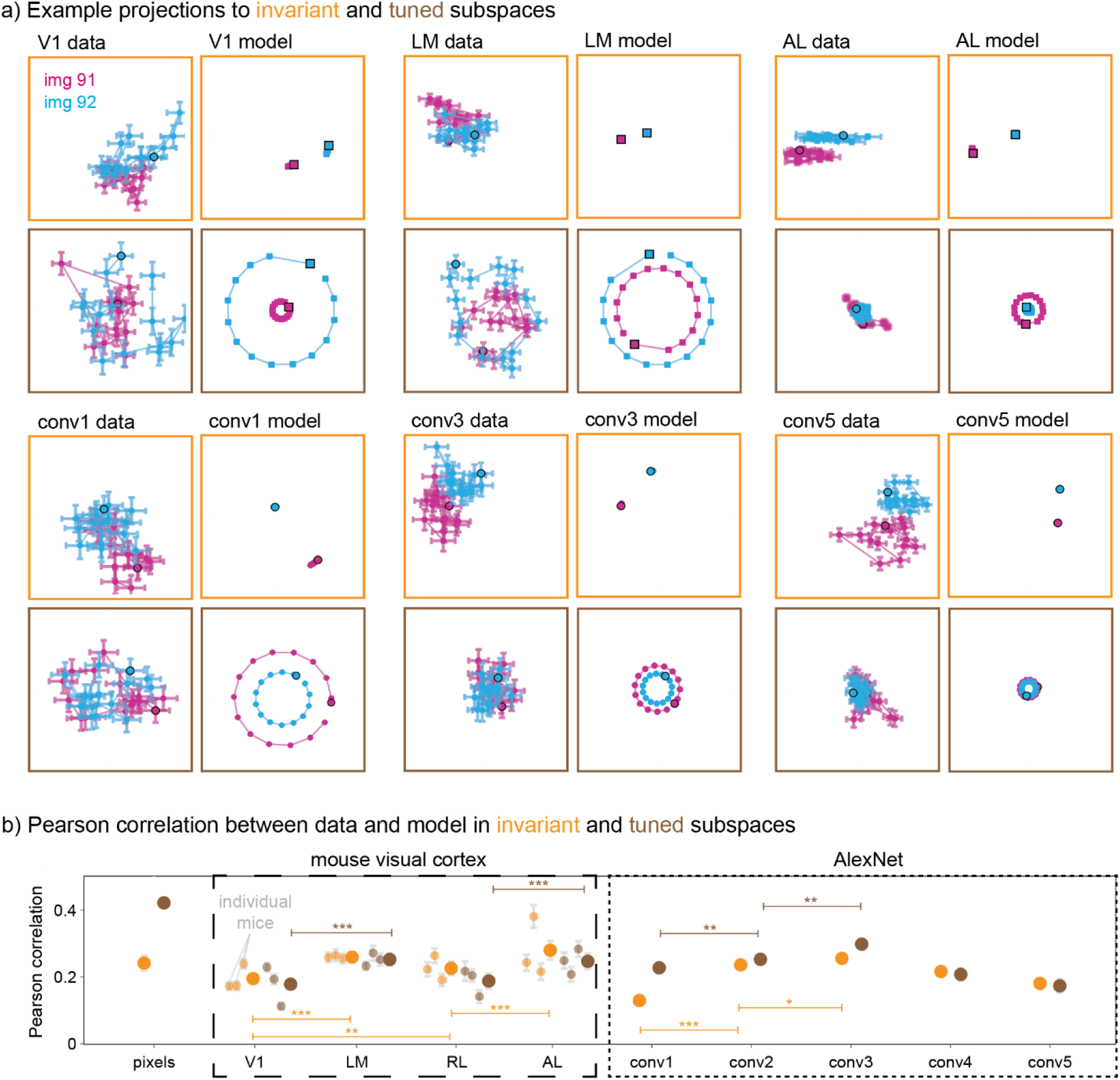
Reshaping of object manifolds within invariant and tuned subspaces differs between mouse visual cortex and AlexNet. **a)** Example projections of held-out neuronal responses to two-dimensional invariant and tuned subspaces, and predictions thereof (as in Fig 3a). For odd columns, dots are means across 80 trials and error bars are sem’s; for even columns, squares are predictions from the equivariance model. We chose to visualize the two dimensions with eigenphases closest to integer multiples of Δϕ. All axes are centered around the estimated rotation center *c* and have the same range for all 4 panels in each area. We used *d* = 20, *N* = 250 random neurons, and mouse data is from mouse 2. **b)** Pearson correlation between held-out data and equivariance model within the invariant and tuned subspaces. For either subspace, we took the mean Pearson correlation across all corresponding dimensions. Shown are mean ± sem across 46 samples, with 3 times more samples for mouse areas. Invisible error bars are smaller than dot sizes.

Projections to the invariant subspace showed a clear separation between held-out identities, which increased across areas. In the invariant subspace, variability with orientation was substantial but differed across areas. The degree of invariance was quantified by the Pearson correlation between data and model in the invariant subspace (averaged across all invariant dimensions). In mouse visual cortex, this Pearson correlation was larger in LM and AL than in V1 and RL, while in AlexNet, it gradually increased along conv1-3 but then decreased slightly in conv4-5 (**Fig 4b**).

Projections to the tuned subspace displayed loops that were more circular and concentric in some areas than others. The degree of tuning (or circularity) was captured by the Pearson correlation between data and model in the tuned subspace (averaged across all tuned dimensions), where the model predicts concentric circles. The Pearson correlation was larger in LM and AL than in V1 and RL, and it gradually increased along conv1-3, but then it dropped in conv4-5.

Our analysis reveals that classification performance under approximate equivariance depends on three distinct geometric features: the dimensionality of the invariant subspace, the degree of invariance, and untangling. Relative orientation decoding depends instead on the existence of the tuned subspace and on the degree of tuning. Pixels show that these geometric features are independent: the degree of invariance in pixels is moderate, yet classification is chance-level because the invariant subspace encodes low-level image statistics that do not linearly separate by image identity. Conversely, the degree of tuning in pixels is the highest of any system studied, with the lowest relative orientation estimation error, because image rotations correspond to exact geometric transformations in pixel space.

Mouse visual cortex maintains invariant and tuned subspaces of fixed dimensionality across the visual hierarchy, with the degrees of invariance and tuning increasing from V1 and RL to LM and AL as object manifolds become more point-like in the invariant subspace and more circular in the tuned subspace (**Fig 4b**; **Extended Data Fig 10b-c**). This combination accounts for the simultaneous improvement in classification and relative orientation decoding observed across areas. AlexNet conv1-3 shows similar geometric structure to mouse cortex; conv4-5 takes a different route, expanding the dimensionality of the invariant subspace at the cost of the degree of tuning. Whether this late-stage organization corresponds to mouse visual cortical areas not recorded here remains an open question.

## DISCUSSION

By combining untangling and equivariance into a common geometric framework, we reconciled two major theories of invariant object classification across vision neuroscience and computer vision. Moreover, we extended them beyond classification to show that object manifolds equivariant to in-plane rotations are structured into orthogonal invariant and tuned subspaces—enabling simple decoding of both object identity and relative orientation. This geometric structure offers a unified solution to the competing demands of invariance and tuning in object recognition.

Equivariance is not a structure that visual processing must construct de novo—it is inherited from the geometry of the input, and what changes across the visual hierarchy is how this inherited structure is reorganized. Consistent with this, invariant and tuned subspaces persisted across all recorded mouse visual cortical areas, but object manifolds were reshaped: they were more point-like within the invariant subspaces of LM and AL compared to V1 and RL, and more circular in the tuned subspace. This suggests parallel untangling along both the putative ventral (V1 → LM) and dorsal (V1 → RL → AL) streams.^36,39–41^ However, a limitation of our experimental setup is that the monitor in front of the mouse only covered about 70° of the vertical visual field, which failed to excite many visual neurons, especially in RL.^56,57^ As a consequence, retinotopy and our derived area boundaries were imprecise. Furthermore, future experiments involving other higher-order visual cortical areas (e.g. AM, POR) are needed to test if these trends hold up at later stages.^36,39–41^

Decoding performances with *N* = 250 neurons are limited by subsampling inherent to cross-area comparisons; ranking is preserved across *N*, and at maximum *N* V1 and LM converge, suggesting compression in LM. We expect this convergence to split for richer image transformations, revealing further differences across visual cortical areas. Even with planar rotations alone, the above-chance linear classification in V1 suggests that untangling starts before visual cortex. The untangling literature^1–4^ has typically located untangling within the primate ventral stream because Gabor models of V1 have near chance-level classification performance, like pixels. Our V1 data show that linear separability is already present at this stage.

A key question in population coding is whether multiple task-relevant variables are encoded by segregated neuronal subpopulations or by a joint, mixed-selective population. Our decoder weight analyses directly address this: identity and relative orientation were decoded from overlapping neurons, providing evidence that mouse visual cortical areas employ a joint rather than a segregated representation. This aligns with the growing view that mixed selectivity is a fundamental feature of cortical population codes and helps with linear separability.^48–50^ Our equivariance framework characterizes a geometrically structured component of the population code (**Extended Data Fig 10b**), describing how such mixed representations are organized: into orthogonal subspaces that enable downstream neurons to independently access both variables. This component, while not capturing all of the response variance (**Extended Data Fig 10c**), contains the dominant geometric structure relevant for decoding image identity and relative orientation (**Fig 3c**). Our framework is agnostic to the specific circuit mechanisms that generate this geometry. Prior work on equivariant neural networks propose biologically plausible learning algorithms that could give rise to such representations.^58,59^ Identifying the biological implementation in mouse visual cortex is an important direction for future work.

The experimental paradigm—naturalistic images presented under rapid serial presentation—is clearly simplified relative to the full complexity of natural visual experience. Nevertheless, this paradigm has been widely used for probing visual representations in both primate and rodent visual cortices and provided fundamental insights into the neuronal basis of object recognition.^1–3,33,44^ In primates, neuronal responses measured under this paradigm have been shown to closely match those in behavioral contexts.^43^ In mice, there is strong behavioral evidence that they can discriminate stimuli similar to the ones used here.^14,15,37^ The absence of a behavioral task reflects our goal of characterizing fast visual processing of as many visual stimuli as possible. We interpret our results as reflecting structure of the population code in mouse visual cortex, while recognizing that future experiments in behaving animals will be important to assess how this structure is used during active vision.

The comparison between mouse visual cortex and AlexNet trained on image classification revealed both agreement and divergence in how visual systems balance invariance and tuning. Our goal with this comparison was not to use the artificial neural network to model neuronal responses in visual cortex, which has been quite successful^23,27^ especially upon adding equivariance-inspired constraints.^29,30^ Rather, we used AlexNet as a controlled visual system in which equivariant structure can be characterized and compared across processing stages. We found that AlexNet conv1–3 showed equivariant structure comparable to that of mouse visual cortex, while conv4–5 achieved further untangling by expanding the dimensionality of the invariant subspace at the expense of tuning. Whether conv4–5 corresponds to other mouse areas not recorded here is an open question.

We analyzed four additional network variants—trained on relative orientation estimation, joint classification and relative orientation estimation (multi-task), classification with rotational augmentation, and with random initialization. The randomly initialized network, which shares the identical architecture as the default classification network but shows no invariant expansion across layers, demonstrates that the expansion of invariant dimensions observed in trained networks is task-driven rather than a consequence of architectural features such as growing receptive field sizes. Whether incorporating richer image transformations (e.g. slant rotations) would constrain this expansion remains an open question for future work.

More broadly, our findings establish equivariance as an organizing principle of neuronal population codes—one that naturally reconciles tuning and invariance within a single geometric framework. While we focused on in-plane rotations in visual cortex, the same framework may apply to other transformations and other brain areas that represent spatial relationships, such as for spatial navigation and motor planning. We hope that this geometric perspective will provide a new lens for understanding how the brain encodes not just what is in the world, but how it changes.

## ACKNOWLEDGEMENTS

This research was funded by the Howard Hughes Medical Institute (HHMI) at the Janelia Research Campus. We thank the Vivarium staff for animal husbandry, Sarah Lindo and Sal DiLisio for surgery support, Jon Arnold for designing headbars and coverslips, Dan Flickinger for microscopy support, and Tobias Goulet for engineering support. We further thank Ann Hermundstad, Arthur Zhao, and Michael Reiser for critical reading of the manuscript, Michael Bonner for suggesting the representational similarity analysis, and Carsen Stringer for fruitful discussions.

## DATA AND CODE AVAILABILITY

The data from the mouse visual recordings is available on https://doi.org/10.25378/janelia.29955530. The code to analyze this data and generate figures can be found on https://github.com/romani-lab/Equivariance-in-neural-networks.

## ONLINE METHODS

### Experimental methods

All of our experiments comply with the standards of the Institutional Animal Care and Use Committee (IACUC) at HHMI Janelia.

#### Animals

We used a total of 3 mice. These mice were bred to express GCaMP6s in excitatory neurons (the particular strain is TetO-GCaMP6s x Emx1-IRES-Cre). All 3 mice were male and about 10 months old. They were housed with a reverse light cycle, and their cages contained a running wheel.

#### Surgical procedures

Surgeries were performed in adult mice (P85 for mouse 1 and 2, P184 for mouse 3), as described in Stringer et al (2021).^47^ In short, mice were anesthetized with Isoflurane before a craniotomy was performed. Marcaine (up to 8 mg/kg) was injected subcutaneously beneath the incision area. Warmed fluids with +5% dextrose and Buprenorphine (0.1 mg/kg, a systemic analgesic) were administered subcutaneously, along with Dexamethasone (2 mg/kg) which was given intramuscularly. The bregma-lambda distance and the location for a 4 mm circular window over V1 were measured, positioning it as far lateral and caudal as possible without compromising implant stability. A 4+5 mm double window was placed into the craniotomy, with the 4 mm window replacing the removed bone piece and the 5 mm window overlapping the bone edge. After surgery, Ketoprofen (5 mg/kg) was administered subcutaneously, and the mice were allowed to recover on heat. They were monitored for pain or distress, with Ketoprofen (5 mg/kg) given for two days following surgery.

#### Imaging data acquisition

We imaged neuronal activity using a custom-built dual-plane 2-photon mesoscope^61^ and employed ScanImage^62^ for data acquisition. The two planes were about 200 and 250 μ*m* below the brain’s surface (in layer 2/3). To correct for Z and XY drift during recording, we used a custom online module, which is now integrated into ScanImage. Following the method described in Stringer et al (2021),^47^ we upgraded the mesoscope to enable temporal multiplexing^10^, approximately doubling the number of recorded neurons. The mice could run freely on an air-floating ball and were acclimated to running on the ball for several sessions before imaging.

#### Stimuli

Half of our 92 different naturalistic images were randomly chosen from the COCO^63^ dataset (odd img) and the other half were naturalistic textures (even img; examples shown in **Extended Data Fig 5a** top). The COCO dataset mainly consists of scenes and objects (with 91 categories; of which we randomly chose 1 image each from 46 categories). The naturalistic textures were either photographed by us or taken from the internet (copyright-free). We first converted the 92 RGB images to grayscale, cropped them into square format (by cutting the longer dimension to the shorter one, starting from the top-left), and resized them to 200×200 pixels. Then we applied a circular mask (gray outside a circle of radius 60 pixels) and histogram-equalized the masked images using the MATLAB package SHINE.^64^ After creating 15 rotated versions of each image (using the MATLAB function ‘imrotate’ with the angles 2π*k*/*n* for *k* = 0, …, *n* − 1 and *n* = 15, and using the parameters ’bicubic’ and ’crop’), we took the central 120×160 pixels and resized the image to 300×400 pixels.

#### Stimulus presentation setup

Mice were head-fixed on an air-suspended, spherical treadmill, on which they could run at will, but this had no consequences on the stimulus presentation. They were surrounded by 3 monitors (one in front, one to the left at right angles, and one to the right at right angles), which were all at about 11 cm from the center of the spherical treadmill. Each monitor is about 16.5×22 cm^2^ (aspect ratio of 4:3). The front monitor spans about 70° × 90°. To prevent direct contamination of the photomultiplier tube from the screen, we placed gel filters in front of each monitor to block green light, transmitting only light from the blue and red channels. We also inserted a Fresnel lens in front of each monitor to slightly magnify images and reduce corner effects between the front and side monitors.

We followed the experimental paradigm of measuring neuronal responses while randomly flashing images in quick succession. The stimulus frame rate was synchronized to the imaging acquisition frame rate at 3.16 Hz. For stimulus presentations, we randomly picked the front or left monitor and one of the 92×15=1380 images, making sure that each image of img 1-90 was shown twice (“2x” images) and each image of img 91-92 eighty times (“80x” images). Each image presentation was interleaved by one gray screen (also shown for about 300 ms). After a block of 300 frames, we inserted a prolonged gray screen period of 20 frames, which we used to estimate baseline neuronal responses. The specific random sequence of monitor choices and images was the same in all recordings.

We used the library PsychToolbox-3^65–67^ in MATLAB to handle the stimulus presentation during the experiment. One frame before each image presentation (during the 1-frame gray screen), we recorded the mouse’s pupil position (using a customized version of Facemap^68^); if it had moved horizontally, we would move the image on the monitor by the corresponding horizontal translation.

### Data analysis methods

#### Processing of calcium imaging data

Calcium imaging data was processed using Suite2p.^11^ It handles motion correction, ROI detection, cell classification, neuropil correction, and spike deconvolution. For details we refer to Stringer et al (2019).^34^ For non-negative deconvolution, we used a decay timescale of 0.75 seconds.^11^

To estimate the neuronal response driven by the images, we also followed a procedure introduced in Stringer et al (2019).^34^ In short, the activity during image-presentation periods for each recorded neuron was z-scored by the baseline activity during the prolonged gray screen periods, which means that we subtracted the mean and divided by the standard deviation. We then orthogonally projected this z-scored quantity (as a vector, with one entry for each neuron) to the subspace that is orthogonal to the baseline PC 1-100, which was computed from the neuronal activity during all prolonged gray screen periods.

Our next preprocessing step, z-scoring neuronal responses across all recorded neurons, reduces multiplicative gain modulations, i.e., a global multiplicative factor at each time point which might arise if, for example, the mouse is running or its pupil area is changing. One indication for global multiplicative gain is if the population response vector at each time has the same mean and standard deviation (see also *Multiplicative gain model*). We note that the resulting population response vector limited to one area no longer has a constant mean and standard deviation. The claim that z-scoring across neurons reduces global gain for small gains is proven in the **Supplementary Information**.

These last two transformations, the removal of baseline activity and global multiplicative gain, made the results cleaner but our main results still held without them (data not shown).

Finally, for the decoders and the equivariance method, we divided preprocessed neuronal responses by the standard deviation across training data for each neuron (“diagonal whitening”). This was to normalize the scale across neurons, as is standard for decoding analyses, and to change to a basis in which *R*(Δϕ) is orthogonal, as derived in the **Supplementary Information**.

#### Retinotopy

Retinotopic maps were computed as described in Zhong et al (2025),^15^ and are shown in **Extended Data Fig 1a**. In short, we compared the output from Suite2p (but only for responses to the 90 training image classes) to simulated (linear) activities from 200 optimized spatial filters of a reference mouse (the filters were obtained from a convolutional model). The most correlated simulated activity determined a spatial position (by the position of the corresponding filter on the image). These spatial positions were then aligned to those of the reference mouse, on which brain areas were outlined following Zhuang et al (2017).^56^

#### Selection of neurons

For our analyses, we selected neurons based on their trial-to-trial correlation to img 1-90 (“2x CC half” in **Extended Data Fig 1b**), which we computed by randomly choosing one single-trial neuronal response for each of img 1-90 and correlating it with the other single-trial neuronal response.^34^ We did this 100 times (over different choices of single trials) and their mean gave our estimated 2x CC half. The selected neurons had a 2x CC half>0.1; corresponding neurons are referred to as “reliably responsive”. A similar computation gave the “80x CC half” for img 91-92 but in this case we randomly chose (also 100 times) forty trials for each image and correlated their means (each mean was over different forty trials).

For most analyses, we randomly sampled 250 reliably responsive neurons in each area without replacement, for at most 100 random samples. For instance, for pixels we used 100 random samples because it contained more than 25,000 units but for AL, we only took 1 sample as it contained less than 500 neurons. In all areas with more than 1 sample, we computed the mean over as many samples without replacement as possible.

In the main text, we only analyzed neuronal responses to the front monitor because for the left monitor the number of reliably responsive neurons in some of the higher-order areas is small (<120, see **Extended Data Fig 1b** bottom). The only areas for which we had a good coverage of reliably responsive neurons for images shown on the left were V1 and the higher-order area PM (posteromedial). Main results for the left monitor are shown in **Supplementary Fig 5**. The main results are consistent for responses to front and left monitors in V1, and we also observe untangling and equivariance in V1 → PM.

#### Single neuron properties

For each reliably responsive neuron, we computed the sensitivity index (“d prime”) and the orientation selectivity index (“OSI”), which capture how much single neuronal responses differ across image classes and across orientations, respectively. For the “2x d prime”, for each pair of img 1-90, we computed the absolute value of the difference in trial-averaged mean (across orientations) responses and divided it by the root mean square of the standard deviation. Then we took the mean across unequal pairs. A similar calculation gave the “80x d prime” for img 91-92 which had only one unequal pair of image classes.

For the OSI, for each neuron and for each image class, we first found the orientation θ_*p*_ where the trial-averaged neuronal response was largest; we call this the “preferred” response. Then we computed the “nonpreferred” response as the mean response at the orientation θ_*p*_ circularly shifted by 168° and 192° (since we do not have θ_*p*_ shifted by 180° in our data). As the sum of preferred and nonpreferred responses could be close to 0, we subtracted from both the minimum response across the 15 orientations (which was always nonzero in our case). Then the difference between the preferred and nonpreferred response divided by their sum gave the OSI for each image class. We took the mean over either img 1-90 or img 91-92 to get one “2x OSI” and one “80x OSI” for each neuron.

Distributions of CC half, OSI and d prime across neurons in each area are shown in **Extended Data Fig 1b** bottom, while values for some example neurons are given in **Supplementary Fig 1**.

#### Class decoder

In the simplest case of two classes, a linear support vector machine (SVM) finds the best hyperplane that separates the two classes of points (in possibly high dimensions). We use a “soft-margin” which allows for imperfect training. The width of the classification margin is controlled by one parameter, typically denoted *C*, which we set to the value *C* = 0.001 (Fig 1d) or *C* = 1.0 (Fig 3c); results for different *C* are shown in **Extended Data Fig 3a** and **Supplementary Fig 4**, respectively. For multiple classes, the SVM performs multiple “one-vs-rest” binary classifications (one per class). The class of a new point is then determined by the binary classifier with the highest confidence score, which is proportional to the signed distance of the point to the corresponding hyperplane. The accuracy *Acc* of classifying one point is either 0 (wrong) or 1 (correct), and the classification error *Err* equals 1 − *Acc*.

We trained a linear SVM on two single-trial neuronal responses to all 92 image classes at 14 consecutive orientations (for img 91-92, we selected two out of the eighty trials). We tested this SVM on trial-averaged neuronal responses to all 92 image classes at the single held-out orientation. We did this 15 times such that each orientation was tested once. We had 92 × 15 = 1380 held-out samples.

The SVM was implemented in python with the scikit-learn library ‘sklearn’ and ‘svm.LinearSVC’. We did not apply any other normalization to the data (e.g. we did not use the ‘preprocessing.StandardScaler’ function).

#### Orientation decoder

To decode the relative orientation angle for neuronal responses to the same image at two different orientations, we first projected neuronal responses to a fixed shared plane and then simply compared the angle between two projected neuronal responses as the estimated relative orientation angle.

The shared plane was defined from trial-averaged neuronal responses to 91 image classes at all orientations (for the trial-average, we used two trials in all cases). Concretely, for each image class, we subtracted the mean response across orientations, and then computed the PC 1-2 from the mean-subtracted responses to the 91 image classes, which defined the two axes of the shared plane. For the held-out image class, instead of mean-subtraction within class, we subtracted the mean response to the 91 image classes at all orientations. After projecting the 15 held-out trial-averaged, subtracted responses to the previously found PC 1-2, we computed the angles between projected responses for orientations ϕ > 0° and ϕ = 0°, resulting in 14 predictions of orientation differences, Δϕ̂. We did this 92 times such that each image class was held-out once. We had 14 × 92 = 1288 held-out samples.

### Computational models

#### Pixel model

For the pixel model, we first cropped the central 300×300 pixels of the 300×400 images that we used for the mouse experiments. We then transformed these 300×300 pixels the same way that images for AlexNet are typically transformed, i.e., we converted them to RGB (using ‘skimage.color.gray2rgb’ in python), rescaled them to 256×256, cropped the central 224×224 pixels, set their means to (0.485, 0.456, 0.406) and their standard deviations to (0.229, 0.224, 0.225) for the 3 RGB channels, respectively. Finally, we took the first (red) channel activations to end up with artificial “neuronal responses” of 224×224=50176 pixels. To each response, we added noise as described in the section *Gaussian noise model* or *Multiplicative gain model*.

Artificial neurons were selected as described in *Selection of neurons*, resulting in 49607 reliably responsive artificial neurons. Like for biological neurons, we divided artificial neuronal responses by the standard deviation across training data for each neuron before decoding and applying the equivariance method.

#### Default AlexNet model (classification only)

To access AlexNet pretrained on ImageNet-1k, we used the library ‘torchvision’ in python. The model was loaded using ‘models.alexnet’ with the parameter ‘weights=DEFAULT’. We used the library ‘torch_intermediate_layer_getter’^69^ to obtain the units’ activities in the convolutional layers.

We downsized the images such that filters of convolutional units in conv 1 (which are of size 11×11) cover approximately the same image area as the receptive fields in V1 (which span about 20 degrees of visual angle^34^). For this, we scaled the images to size 40×40 and embedded them in the center of a gray image of size 256×256. We then further transformed these images as described in the section *Pixel model*. Addition of noise and selection of artificial neurons also proceeded as for pixels. The number of reliably responsive artificial neurons was 5045 in conv1, 2363 in conv2, 938 in conv3, 551 in conv4, and 309 in conv5.

#### Gaussian noise model

For a more direct comparison between biological and artificial neuronal responses, we added Gaussian noise to artificial neuronal responses to simulate trial-to-trial variability. For each artificial neuron, we sampled from a Gaussian distribution of mean 0 and standard deviation equal to 1.98 times the neuron’s standard deviation across img 1-90 (all orientations). The factor of 1.98 is the estimated square root of the noise-to-signal power ratio from all reliably responsive neurons in mouse 1, i.e., the mean of 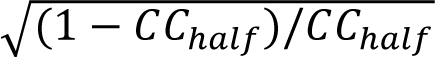 where *CC*_*half*_ is the “2x CC half” introduced in the section *Selection of neurons* (this equation can be obtained from Ref. [70], combining Eq. (9) and (17) for N=2 trials). We chose Gaussian noise for simplicity, using it as a coarse approximation—not to model individual response variability in detail, but to ensure that the same decoding and subspace analyses could be applied identically to both artificial and biological neuronal responses.

#### Multiplicative gain model

To simulate biologically realistic trial-to-trial variability, we also implemented a multiplicative gain model (**Extended Data Fig 7**). As claimed in the section *Processing of calcium imaging data*, we first verified that z-scoring responses largely removed noise correlations between neuron pairs (panel a), suggesting that shared variability is primarily multiplicative in nature. Consistently, the standard deviation of neural responses scaled approximately linearly with the mean in the raw data, but this relationship was removed after z-scoring responses across neurons (panel b), confirming that shared variability is primarily multiplicative in nature. A single shared gain component captured about 17% (6%) of the total trial-to-trial variance for the 2x (80x) stimuli (panel c).

Following Ref. [60], we model the activity of neuron *i* at trial *t* with stimulus *s*_*t*_ as *r*_*i*_ (*t*) = *f*_*i*_ (*s*_*t*_)(1 + *g*_*t*_α_*i*_), where the mean response of neuron *i* to stimulus *s*_*t*_ is given by the parameter *f_i_*(*s*) = 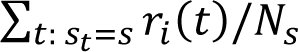 with *N*_*s*_ being the number of trials *t* with *s*_*t*_ = *s*, the trial-to-trial variability is modeled by *g*_*t*_ which satisfies ∑_*t*: *st*=*s*_ *g*_*t*_ = 0 for each stimulus *s*, and the effect of gain on neuron *i* is modeled by α_*i*_ with ∑_*i*_ α_*i*_ ≥ 0 and ∑_*i*_ α_*i*_ ^2^ = 1.

To estimate the parameters *f*_*i*_ (*s*_*t*_), *g*_*t*_, and α_*i*_ of the multiplicative gain model, we performed rank-1 linear regression on the reliably responsive neuronal responses of mouse 1 (6,349 neurons). Specifically, we estimated *f*_*i*_ (*s*) as the mean across trials for each stimulus *s* and for each neuron *i*, computed *r*_*i*_ (*t*)/*f*_*i*_ (*s*_*t*_), subtracted the mean over trials for each *s*_*t*_ = *s*, and then performed rank-1 linear regression on these relative residuals to obtain *g*_*t*_ and α_*i*_ with ∑_*t*: *st*=*s*_ *g*_*t*_ = 0. To impose the remaining conditions, we replaced α_*i*_ by α_*i*_/√∑_*j*_ α_*j*_ ^2^, and if ∑_*i*_ α_*i*_ < 0, we flipped the signs of all α_*i*_ and *g*_*t*_. The fit was performed separately on the 2x and 80x stimuli.

To apply the multiplicative gain model to pixel representations and AlexNet layers, we randomly sampled neuron-specific gain sensitivities α_*i*_ from the empirically estimated distribution, used the same gain fluctuations *g*_*t*_ as in the neural data, and set *f*_*i*_ (*s*) equal to the unit activations of artificial neuron *i* in response to stimulus *s*. Main results (decoding errors, subspace dimensionalities, Pearson correlations, model fit, and variance explained) are shown in panels d-f, which are similar to the main results obtained with additive Gaussian noise (main figures).

#### Additional AlexNet models

We additionally trained three AlexNet variants on ImageNet-1k from random initialization (**Extended Data Fig 8**): a classification-only model with rotation augmentation (panel a), an orientation-only model (panel b), and a multi-task model combining classification and relative orientation estimation (panel c). All three share the same backbone (conv1-5 plus one fully connected layer), training procedure, and image preprocessing, differing only in their task heads and loss functions.

For the rotation-augmented model, the task head is the same as for the default AlexNet (denoted here as the “classification” network), consisting of two fully connected layers (4096 → 4096 and 4096 → 1000). For the orientation-only model, two images that are rotated versions of each other are independently processed through the same (“Siamese”) backbone to produce two embeddings, which are then concatenated. This vector is then passed through two fully connected layers (8192 → 4096 and 4096 → 2) with the two output values representing the sine and cosine of the rotation from the first image to the second image. The multi-task model uses the same Siamese backbone, followed by a Siamese classification head as well as a joint orientation head. Each image pair is classified independently, and their relative orientation is estimated jointly from the concatenated embedding.

For the training, images were preprocessed following standard ImageNet practice: random resized cropping to 224×224 pixels and random horizontal flipping during training; resizing to 256×256 pixels and center cropping during validation. For rotation-augmentation, a single random rotation angle was sampled uniformly for each training image. For relative orientation estimation, two independent random angles ϕ_1_and ϕ_2_ were sampled for each image. Image rotations were applied using bilinear interpolation (using ‘skimage.transform.rotate’ in python). Finally, a circular mask was applied: pixels outside an inscribed circle (with radius of 112 pixels) were set to a uniform gray value, eliminating corner artifacts that would otherwise provide a trivial cue for rotation angle.

Classification was trained with cross-entropy loss over 1000 ImageNet classes. Relative orientation estimation was trained minimizing the mean-squared error between the output (a, b) of the orientation head and (sin(Δϕ), cos(Δϕ)) where Δϕ ≡ ϕ_2_ − ϕ_1_ mod 360° is the true angle between the image pair. For the multi-task network, the loss was simply the sum of the cross-entropy losses for both images and the mean-squared error for the orientation. All models were trained equivalently to the default AlexNet with stochastic gradient descent with momentum 0.9, weight decay 10^−4^, base learning rate 0.01, and batch size 256. The learning rate followed a step schedule, decaying by a factor of 10 every 30 epochs (i.e., at epochs 30 and 60). Models were trained for a total of 90 epochs. No gradient clipping was applied. Weights were initialized randomly (no pretraining).

To assess the final task performance, we evaluated the network output on 100 randomly sampled ImageNet test images, resized to 256×256, center-cropped to 224×224, rotated at 15 evenly spaced orientations, and masked outside the inscribed circle (panel d). The network output classification error was computed as 1 minus the top-1 accuracy evaluated across the 100 images. The network output relative orientation estimation error was computed as min (|Δϕ̂ − Δϕ|, 360° − |Δϕ̂ − Δϕ|), where Δϕ̂ =arctan(b/a) (using the numpy function ‘atan2’) and (*a*, *b*) is the output of the orientation head with the second image being the canonical orientation (not rotated).

To compare the equivariant structure across network variants, we applied the same analyses as in the main text, using the same preprocessing (resized images, Gaussian noise, N = 250 subsampling) and the same 92 images at 15 orientations (panels e-g). Classification and relative orientation estimation errors (panel e), invariant and tuned subspace dimensions (panel f), and Pearson correlations (panel g) were computed as in the main text.

The default classification network achieved low classification error only for unrotated images, while rotational augmentation and multi-task training had similar errors and were largely independent of orientation (panel d top). The orientation and multi-task networks both achieved near-zero orientation estimation error (< 10°) across all relative orientations, whereas the random network predicted a fixed angle that depended on the random initialization (panel d bottom).

At the neuronal population level, the classification error decreased strongly across layers for the classification, rotation-augmented, and multi-task networks, reaching around 0.6 at conv5, whereas classification errors in the orientation and random networks did not go below 0.8 (panel e). The relative orientation estimation error was lowest for the orientation network and decreased across layers to around 25° at conv5; for the classification, rotation-augmented, and multi-task networks it first slightly decreased until conv3 and then increased to near-chance level, remaining lowest in the multi-task network. The number of invariant dimensions increased across conv1-5 for the classification, rotation-augmented, and multi-task networks but remained low for the orientation-trained and random networks (panel f). Notably, the random network, which shares the identical architecture as the classification network but shows no invariant expansion, indicates that the expansion is task-driven rather than a consequence of architectural features. The number of tuned dimensions remained stable between 4 and 8 across all networks. The degree of invariance was similar across all networks (panel g). The degree of tuning was slightly higher for the multi-task network than for the rotation-augmented network, which suggests that training on relative orientation estimation—more than exposure to rotated images—implicitly results in a circular structure of population responses, as predicted by our theory.

Together, these results show that training objectives strongly shape the equivariant structure across layers in interpretable ways, and that only networks including classification in their training objective show the simultaneous improvement in both classification and relative orientation decoding in early-to-intermediate layers, consistent with results in mouse visual cortex.

### Computational method for equivariance

Previous methods for estimating equivariance were specific to convolutional neural networks.^54^ We developed a method that applies to any set of neuronal responses, be it from artificial or biological visual systems.

#### Estimation procedure

We denote the neuronal population response as an *m*-dimensional vector *r*(ϕ) that depends on an image and its orientation ϕ. Rotation-equivariance means that the response to the image at orientation ϕ + Δϕ is *r*(ϕ + Δϕ) − *c* = *R*(Δϕ) · (*r*(ϕ) − *c*) where · denotes matrix-vector multiplication and *c* is an *m*-dimensional vector—the rotation center. We assume that just like ordinary rotations in space, the *m* × *m* matrix *R*(Δϕ) should be orthogonal, independent of image identity and of its orientation ϕ, periodic in Δϕ with period 360°, and be compositional, i.e., *R*(Δϕ)^*k*^ = *R*(*k*Δϕ) for any integer *k*.

For what follows, it will be more convenient to work with the offset *e*(Δϕ) = *R*(Δϕ) · *c* − *c* instead of *c*, which satisfies *r*(Δϕ) = *R*(Δϕ) · *r*(0) − *e*(Δϕ). Let us denote estimated quantities with a hat. The goal of the estimation procedure is to find both *R̂*(Δϕ) and *e*^(Δϕ) from pairs of neuronal responses to images that are related by an image rotation of Δϕ. Naively, one might expect this problem to correspond to the orthogonal Procrustes problem (see more below). However, the main limitation of the well-known solution to the orthogonal Procrustes problem is that the rank of *R̂*(Δϕ) would be limited by the number of pairs *T*, which may be smaller than the number of neurons *m*. This is a problem because orthogonal matrices must have full rank.

To circumvent this problem, we introduce a *d*-dimensional orthogonal projection such that *R̂*(Δϕ) is estimated in *d*-dimensional space. The projection is accomplished by a *m* × *d* matrix W(Δϕ) which satisfies that W(Δϕ)^*T*^W(Δϕ) is the *d* × *d* identity matrix where the superscript *T* indicates matrix transposition. The previous relationship between neuronal responses then becomes *r*(Δϕ) = W(Δϕ)*R̂*(Δϕ)W(Δϕ)^*T*^ · *r*(0) − W(Δϕ) · *e*^(Δϕ). To lighten notation, we shall drop the explicit Δϕ-dependence in the following except where necessary.

To formalize the goal of our estimation procedure, let us concatenate unrotated responses into a *m* × *T* matrix *A* = (*r*_1_(0), *r*_2_(0), …, *r*_*T*_(0)) and rotated responses into another *m* × *T* matrix *B* = (*r*_1_(Δϕ), *r*_2_(Δϕ), …, *r*_*T*_(Δϕ)). Then the relationship between neuronal responses is simply *B* = W(*R̂*W^*T*^*A* − W^*T*^*Ê*) where *Ê* is a *m* × *T* matrix whose columns all equal *e*^. Our goal is thus to minimize ‖*B* − W(*R̂*W^*T*^*A* − W^*T*^*Ê*)‖_*F*_ for W, *R̂* and *Ê* with their respective constraints, i.e., W is a *m* × *d* matrix such that W^*T*^W is the identity matrix, *R̂* is a *d* × *d* orthogonal matrix, and *Ê* is a *m* × *T* matrix of rank 1.

Here, ‖. ‖ denotes the Frobenius matrix norm which is defined as 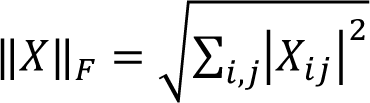 where X_*ij*_ are the matrix elements of a matrix X.

The case of *d* = *m*, where W is the identity matrix, and without the rank-1 constraint on *Ê*, corresponds to the orthogonal Procrustes problem. The well-known solution is as follows.^53^ The orthogonal matrix *R̂* is given by the polar decomposition of *BA*^*T*^, i.e., if the singular value decomposition (SVD) of *BA*^*T*^ is *USV*^*T*^, where *S* is the diagonal matrix with singular values and *U*, *V* are orthogonal, then *R̂* = *UV*^*T*^. The remainder *Ê* is simply *Ê* = *B* − *R̂A*. This solution is unique if the nonzero singular values are nondegenerate (which is generically the case).

If instead *d* < *m*, we note that the best low-rank approximation of *BA*^*T*^is *U*_*d*_*S*_*d*_*V*_*d*_^*T*^ where the subscript means that we only use the first *d* columns (and rows, for *S*_*d*_); this is known as the Eckart–Young–Mirsky theorem. If *A*, *B* are “whitened”, i.e., their covariances equal the identity matrix, then the columns of *U*_*d*_ and *V*_*d*_ correspond to the first *d* pairs of canonical directions. On the other hand, if *A*, *B* are nearly the same (as would be the case for Δϕ ≈ 0°), then *U*_*d*_, *V*_*d*_ are nearly the same and approximately equal the first *d* PCs. As we only allow for one projection, we chose W = *U*_*d*_ (we could have chosen W = *V*_*d*_ instead). After the *d*-dimensional projection, we apply the well-known orthogonal Procrustes solution to obtain estimates of *R̂* and *Ê*. Given W and *R̂*, one can show that the W^*T*^*Ê* that best achieves the estimation goal with the rank-1 constraint is given by the mean of W^*T*^*B* − *R̂*W^*T*^*A* across columns.

In summary, our estimation procedure is as follows:

1. Compute the SVD of *BA*^*T*^ = *USV*^*T*^.
2. Define W as the first *d* columns of *U*.
3. Compute the SVD of W^*T*^*BA*^*T*^W = *U*′*S*′*V*′^*T*^.
4. Define *R̂* = *U*′*V*′^*T*^.
5. Define *Ê* as W times the mean of W^*T*^*B* − *R̂*W^*T*^*A* across columns.

We applied the estimation procedure on trial-averaged neuronal responses to 90 image classes (all orientations; two trials). As held-out class pairs, we chose one image from the COCO dataset and one from the texture dataset; specifically, we used the 46 pairs (img 2i − 1, img 2i), resulting in 46 held-out class pair samples. We emphasize that, for each held-out class pair, we only estimated one W, one *R̂*(Δϕ), and one *e*^(Δϕ).

#### Requirements for rotation-equivariance

Successful estimation of W, *R̂* and *Ê* does not guarantee rotation-equivariance. Here we discuss the additional requirements for equivariance.

First, rotation-equivariance imposes 3 conditions on *R̂*: orthogonality, periodicity, and compositionality. Orthogonality is guaranteed by our estimation procedure, but periodicity and compositionality are not. As we show in the **Supplementary Information**, periodicity and compositionality imply that the eigenphases Δθ of *R̂* must be integer multiples of the image rotation angle Δϕ. Not all our estimated Δθs satisfy this condition (**Fig 3b** top). However, we notice that the variance explained of the corresponding eigenspace is often larger for Δθ close to integer multiples of Δϕ, suggesting that other eigenspaces have higher trial-to-trial variability. Therefore, our interpretation is that our analyzed neuronal responses are approximately rotation-equivariant (**Fig 2a**) where some dimensions with large variance explained satisfy the equivariance constraints and others with smaller variance explained do not. To be clear, the variance explained was computed on held-out class pairs (46 samples).

#### Equivariance model

The equivariance model allows us to predict how neuronal responses change with orientation, using the estimated projection matrix W, rotation matrix *R̂*(Δϕ), and shift vector *e*^(Δϕ). If the response to an image is *r*(0), the equivariance model predicts the response to the image rotated by 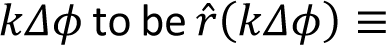 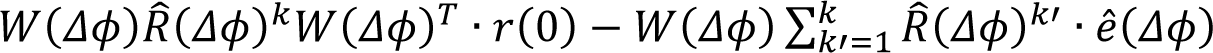 for all integers *k*′ > 0, which assumes compositionality of *R̂*(Δϕ). Therefore, if the equivariance model fits the data well, *R̂*(Δϕ) is likely compositional.

The equivariance model performance can be quantified by the Pearson correlation between data and model in the projected subspace. In **Fig 4b**, we computed the mean Pearson correlation across all dimensions of the invariant subspace (orange) and the mean Pearson correlation across all dimensions of the tuned subspace (brown), so that the two values are directly comparable.

We also assessed the model performance in the full neuronal response space by computing the normalized Pearson correlation, *CC*_*norm*_^70^ (**Extended Data Fig 10b**). This quantity takes into account neuronal trial-to-trial variability. We stress that both the unnormalized Pearson correlation and *CC*_*norm*_ were evaluated on held-out data (46 samples).

#### Decoding in tuned and invariant subspaces

To show that the tuned and invariant subspaces estimated from our computational method capture the trends of the decoding performances in the full neuronal response space (**Fig 1d-e**), we repeated the decoding analyses after projecting neuronal responses to these subspaces (**Fig 3c**). In our computational method, we only used part of the training data of each decoder to find the projections to the tuned and invariant subspaces. Concretely, for the linear SVM, we used the trial-averaged neuronal responses to all 92 image classes at all orientations but one; for the orientation decoder, we used the trial-averaged neuronal responses to 91 image classes (all orientations). This procedure makes sure that held-out data was not used anywhere during training.

### Statistical tests

We performed Wilcoxon signed-rank tests to compare paired samples in **Fig 1d-e**, **Fig 3c**, and **Fig 4b**. Statistical significance was visualized as * if *p* < 0.1, ** if *p* < 0.01, and *** if *p* < 0.001. No adjustments were made for multiple comparisons. In python, we used the function ‘wilcoxon’ of the library ‘scipy.stats’ with the parameter ‘alternative’ equaling either ‘greater’ (for decoding) or ‘less’ (for Pearson correlation).

The exact p-values are as follows:

**- Fig 1d**

mouse visual cortex: 10^−59^ (V1>LM), 1.0 (V1>RL), 1.0 (V1>AL), 1.0 (LM>RL), 10^−25^ (RL>AL)

AlexNet: 10^−103^ (conv1>conv2), 10^−96^ (conv1>conv3), 10^−103^ (conv1>conv4), 10^−55^

(conv1>conv5), 10^−29^ (conv2>conv3), 10^−29^ (conv3>conv4), 0.01 (conv4>conv5)

**- Fig 1e**

mouse visual cortex: 10^−53^ (V1>LM), 0.10 (V1>RL), 10^−10^ (V1>AL), 1.0 (LM>RL), 10^−4^ (RL>AL)

AlexNet: 10^−6^ (conv1>conv2), 10^−26^ (conv1>conv3), 10^−5^ (conv1>conv4), 0.99 (conv1>conv5),

10^−15^ (conv2>conv3), 1.0 (conv3>conv4), 1.0 (conv4>conv5)

**- Fig 3c top**

mouse visual cortex: 10^−46^ (V1>LM), 1.0 (V1>RL), 1.0 (V1>AL), 1.0 (LM>RL), 10^−9^ (RL>AL)

AlexNet: 10^−60^ (conv1>conv2), 10^−81^ (conv1>conv3), 10^−85^ (conv1>conv4), 10^−29^

(conv1>conv5), 10^−25^ (conv2>conv3), 10^−14^ (conv3>conv4), 0.05 (conv4>conv5)

**- Fig 3c bottom**

mouse visual cortex: 10^−62^ (V1>LM), 10^−7^ (V1>RL), 10^−42^ (V1>AL), 1.0 (LM>RL), 10^−14^ (RL>AL)

AlexNet: 10^−18^ (conv1>conv2), 10^−20^ (conv1>conv3), 10^−8^ (conv1>conv4), 1.0 (conv1>conv5),

10^−4^ (conv2>conv3), 0.99 (conv3>conv4), 1.0 (conv4>conv5)

**- Fig 4b invariant (orange)**

mouse visual cortex: 10^−11^ (V1<LM), 0.008 (V1<RL), 10^−6^ (V1<AL), 0.99 (LM<RL), 10^−4^ (RL<AL)

AlexNet: 10^−12^ (conv1<conv2), 10^−14^ (conv1<conv3), 10^−7^ (conv1<conv4), 0.001

(conv1<conv5), 0.03 (conv2<conv3), 0.99 (conv3<conv4), 0.98 (conv4<conv5)

**- Fig 4b tuned (brown)**

mouse visual cortex: 10^−11^ (V1<LM), 0.08 (V1<RL), 10^−6^ (V1<AL), 0.99 (LM<RL), 10^−4^ (RL<AL)

AlexNet: 0.005 (conv1<conv2), 10^−5^ (conv1<conv3), 0.88 (conv1<conv4), 0.99 (conv1<conv5),

0.001 (conv2<conv3), 0.99 (conv3<conv4), 0.95 (conv4<conv5)

### Decoder comparison and effect size quantification

We compared the two decoders between the full space (**Fig 1d-e**) and the subspaces (**Fig 3c**) using paired held-out samples. The difference between the decoding errors in the full space and the subspaces is denoted Δ. If mean(Δ)/std(Δ) > 0 then decoding is consistently better in the subspace, and if mean(Δ)/std(Δ) < 0 then decoding is consistently better in the full space. To further quantify how well an ideal observer could infer from the decoding error if the decoding was performed in the full space or the subspace, we computed the area under the receiver operating characteristic curve (ROC AUC) using the decoding error as the decision variable. ROC AUC values range between 0 and 1.0, with 0.5 indicating chance-level discriminability. Values greater than 0.5 indicate that decoding in the subspace tends to yield lower errors than decoding in the full space, whereas values below 0.5 indicate the opposite.

We give the estimated effect sizes in the format (mean(Δ)/std(Δ), ROC AUC), first for class decoding:

- pixels: (0.43, 0.56), mouse 1 V1: (−0.09, 0.46), mouse 2 V1: (−0.06, 0.47), mouse3 V1: (−0.09, 0.46), mouse 1 LM: (−0.10, 0.46), mouse 2 LM: (−0.11, 0.47), mouse 3 LM: (−0.06, 0.48), mouse 1 RL: (0.02, 0.50), mouse 2 RL: (−0.01, 0.49), mouse 3 RL: (−0.03, 0.48), mouse 1 AL: (−0.03, 0.49), mouse 2 AL: (−0.09, 0.48), mouse 3 AL: (−0.08, 0.48), AlexNet conv1: (0.37, 0.57), conv2: (0.09, 0.50), conv3: (0.03, 0.50), conv4: (−0.08, 0.48), conv5: (−0.08, 0.47).

and second for relative orientation decoding:

- pixels: (0.32, 0.50), mouse 1 V1: (0.39, 0.57), mouse 2 V1: (0.29, 0.55), mouse3 V1: (0.30, 0.56), mouse 1 LM: (0.26, 0.54), mouse 2 LM: (0.20, 0.55), mouse 3 LM: (0.20, 0.54), mouse 1 RL: (0.16, 0.55), mouse 2 RL: (0.19, 0.55), mouse 3 RL: (0.21, 0.55), mouse 1 AL: (0.35, 0.60), mouse 2 AL: (0.21, 0.55), mouse 3 AL: (0.13, 0.53), AlexNet conv1: (0.27, 0.54), conv2: (0.32, 0.56), conv3: (0.05, 0.50), conv4: (0.14, 0.53), conv5: (0.01, 0.50).

---

**Extended Data Fig 1.**
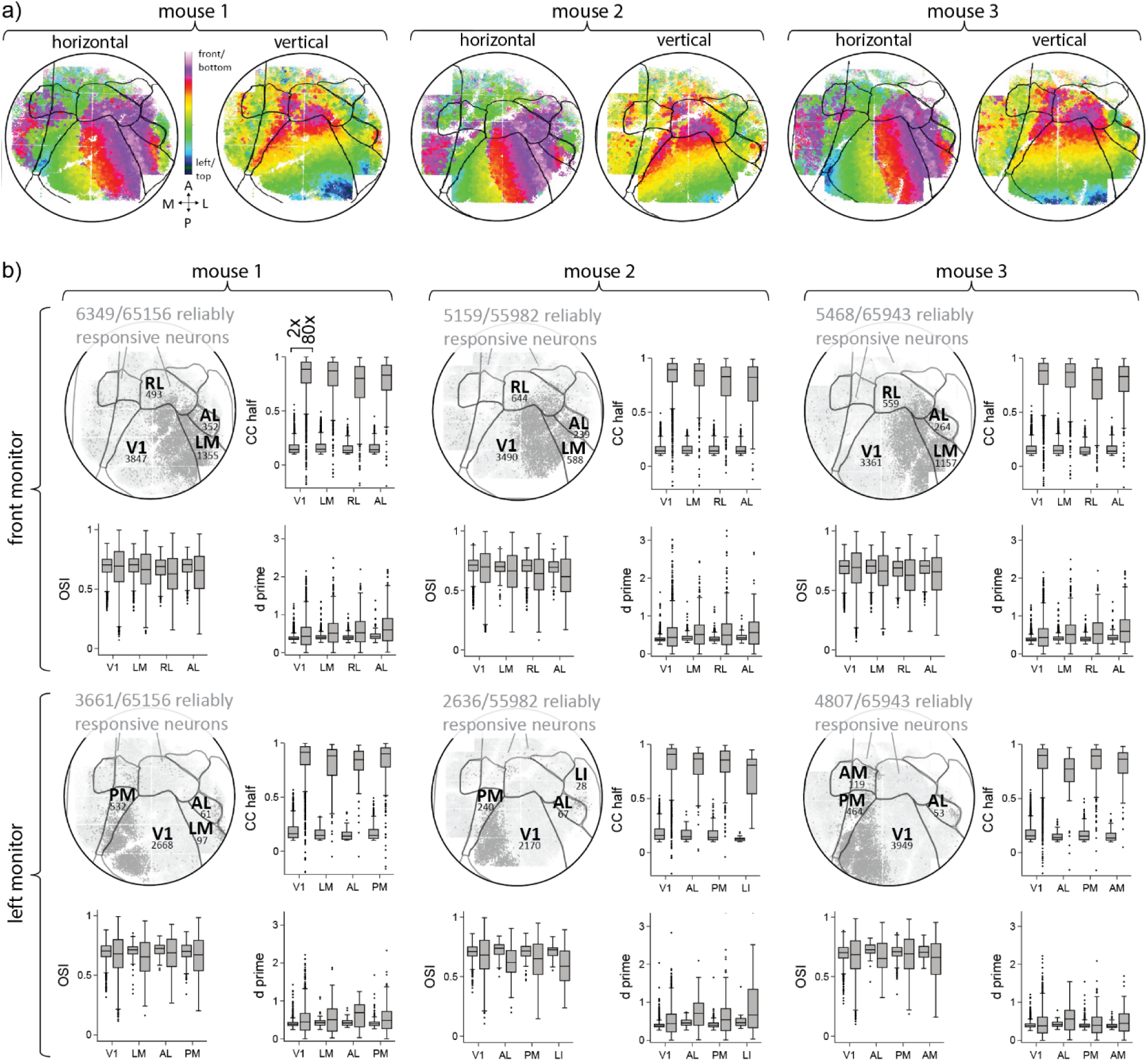
Retinotopy and single neuron distributions across visual cortical areas. **a)** Estimated horizontal (left to front) and vertical (top to bottom) retinotopy in visual cortical areas for the 3 imaged mice. Black lines show estimated area boundaries. **b)** For each mouse, we analyzed neurons that reliably responded to either images on the front (first two rows) or on the left (last two rows) monitor. For each mouse and monitor, the top-left panel shows the reliably responsive neurons in dark gray and all measured neurons in light gray. For each area that we analyzed, we annotated it by its name and the number of reliably responsive neurons it contains. The other 3 panels show single neuron distributions across areas for neuronal responses to 2x and 80x image classes separately, which are the trial-to-trial correlation (“CC half”), the sensitivity index (“d prime”), and the orientation selectivity index (“OSI”). Box plots display the quartiles (horizontal lines of the box) and the interquartile range (IQR; whiskers are 1.5 times the IQR displaced from the first and third quartiles); points are outliers. 2x refers to img 1-90, which were shown twice, and 80x to img 91-92, which were shown eighty times.

**Extended Data Fig 2.**
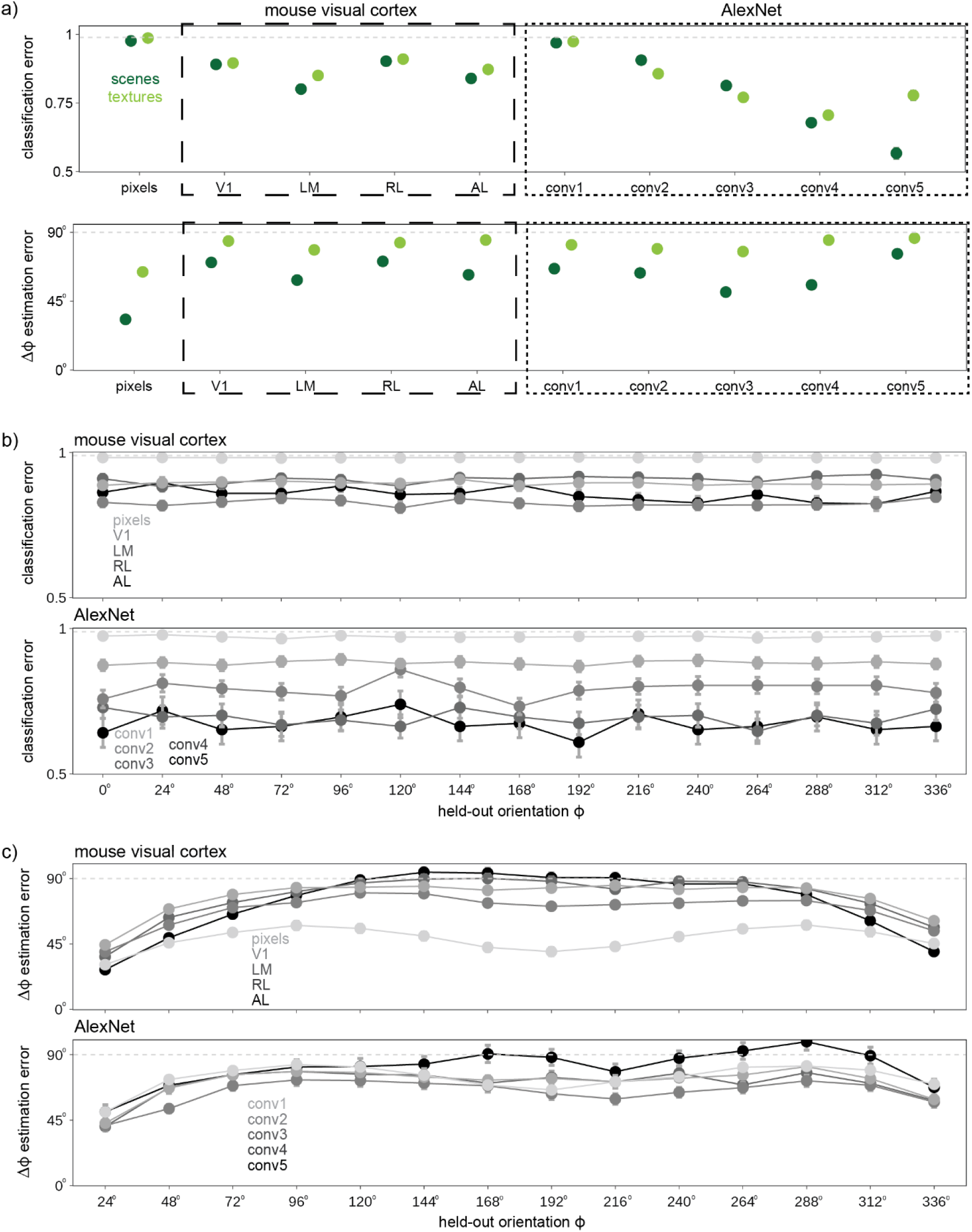
Decoding analyses split up by scenes/textures and orientation. **a)** Analogue of Fig 1d-e but where results are split by scenes (incl. img 91; dark green) and textures (incl. img 92; light green). **b)** Analogue of Fig 1d but results are split by held-out orientation, with pixels and mouse visual cortical areas on top and AlexNet layers at the bottom. **c)** Analogue of Fig 1e but results are split by held-out orientation, with pixels and mouse visual cortical areas on top and AlexNet layers at the bottom.

**Extended Data Fig 3.**
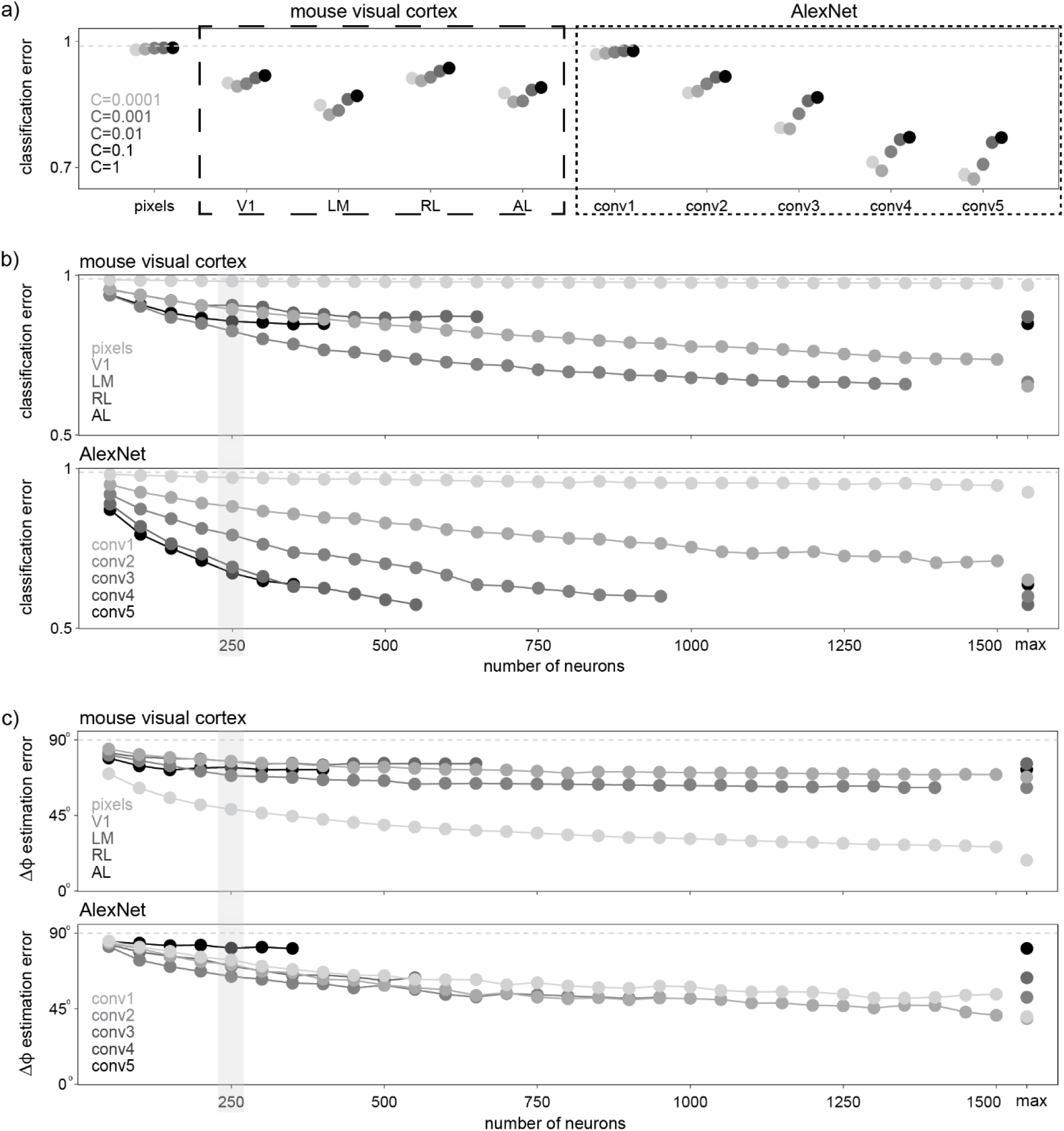
Additional decoding analyses. **a)** Analogue of Fig 1d but for different L2-regularization parameters C used in the linear SVM. **b)** Classification error as a function of numbers of neurons N per area (for *C* = 0.001, as in Fig 1d). N=max indicates that all reliably responsive neurons in an area were used. On top, different shades of gray correspond to pixels and mouse visual cortical areas; on bottom, they correspond to different AlexNet layers. **c)** Similar to b but for the relative orientation decoder. In all panels, results were averaged over at most 100 random samples of *N* = 250 neurons and error bars are smaller than dot sizes. The shaded area in b-c highlights *N* = 250 which we used for most analyses.

**Extended Data Fig 4.**
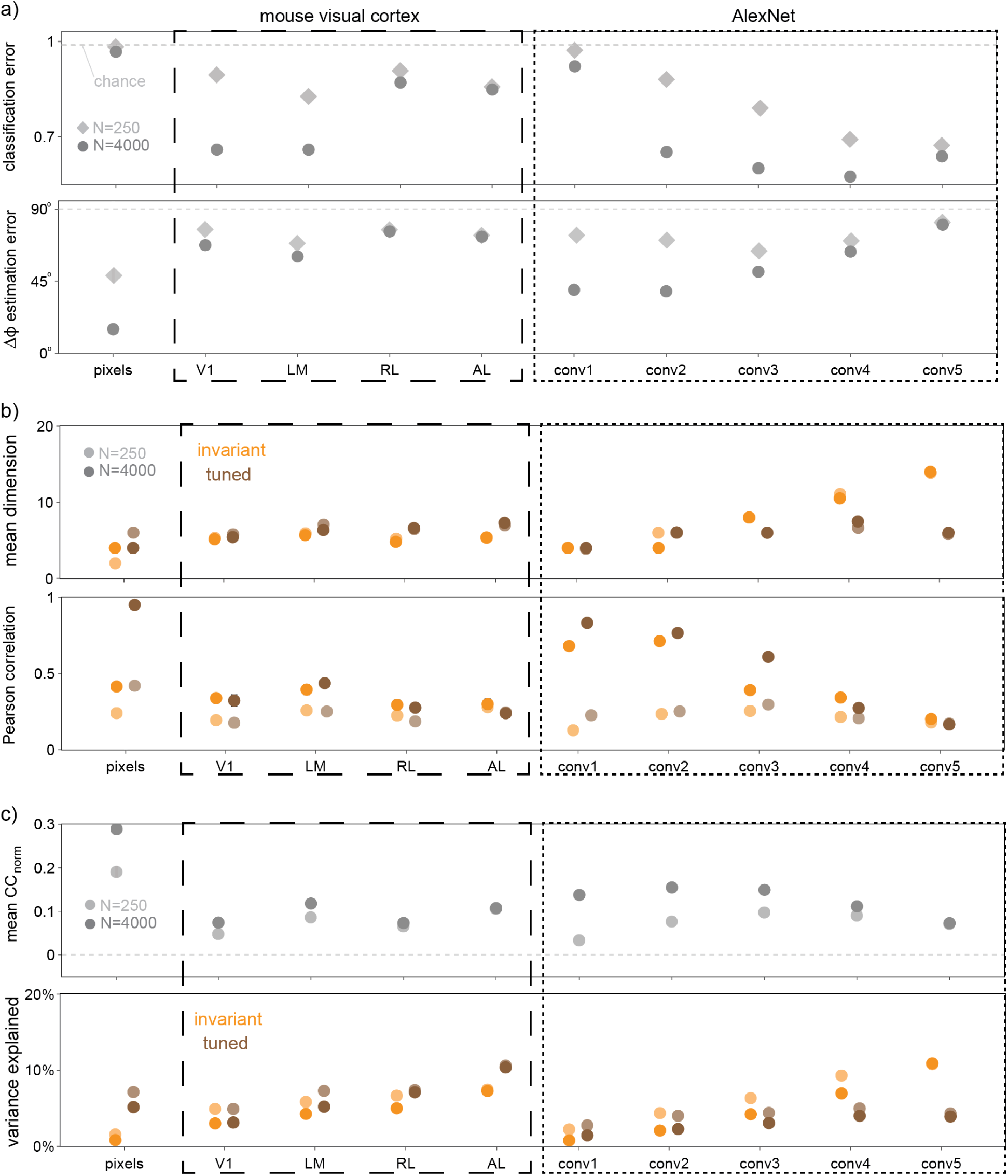
Comparison of main results for N=250 and N=4000 reliably responsive neurons. Analogues of **a)** Fig 1d-e (*C* = 0.001), **b)** Fig 3b bottom and Fig 4b, and **c)** Extended Data Fig 10b-c using up to *N* = 4000 reliably responsive neurons in each area (opaque dots). For comparison, we added the corresponding results for *N* = 250 in semi-transparent.

**Extended Data Fig 5.**
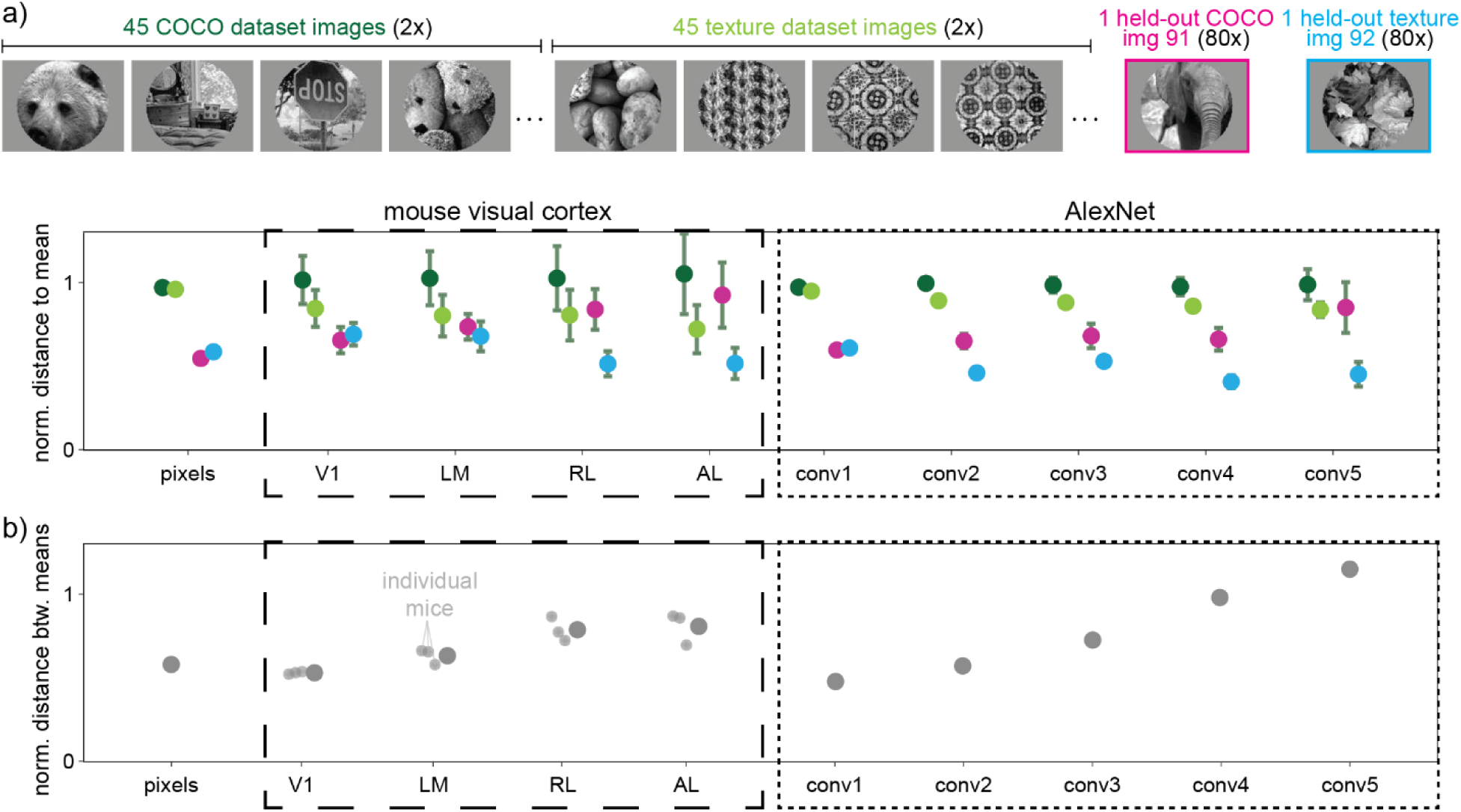
For each image class, neuronal responses are approximately equidistant from mean neuronal response, and distance between mean neuronal responses increases. **a)** Top: Example images from our stimulus set (at reference orientation ϕ = 0°). Images 1-90 were displayed two times (2x) while images 91-92 were shown eighty times (80x) at each orientation. Bottom: Within each area, we computed the Euclidean distance in neuronal response space of each response vector to the mean vector (one per class), normalized by the square root of the dimension (√250). Dots are means and error bars are sem’s over image rotations (675 samples each for the two shades of green and 15 samples for pink and blue). Dark green includes all 45 classes in the COCO dataset (e.g. img 1), bright green all 45 classes in the texture dataset (e.g. img 2), pink is for img 91, and blue for img 92. **b)** Euclidean distance in neuronal response space between the mean vector to img 91 and the mean vector to img 92, normalized by the standard deviation of neuronal responses to img 91 and img 92. Results are averaged over at most 100 independent random samples of *N* = 250 neurons.

**Extended Data Fig 6.**
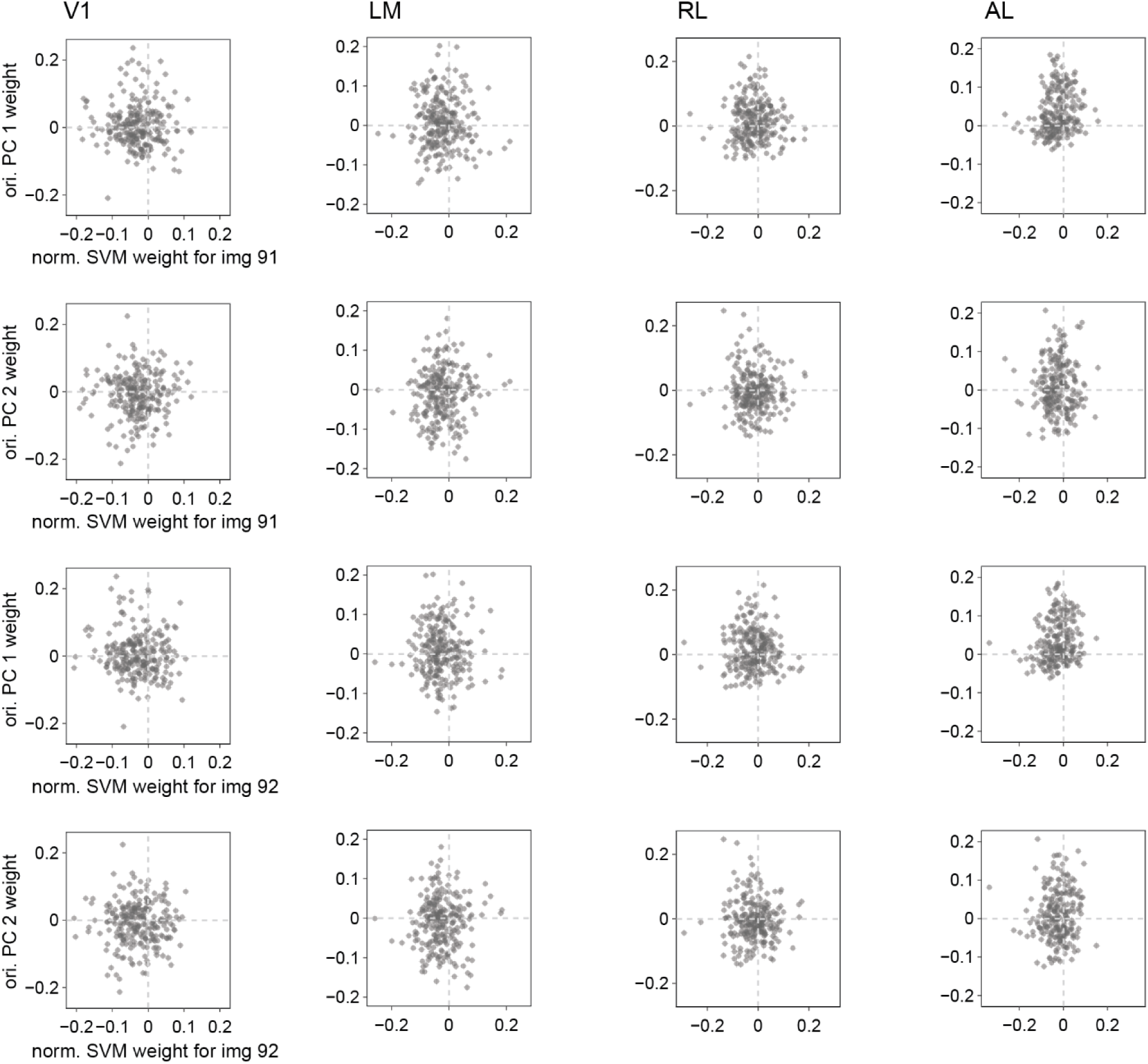
Weights from class decoder and relative orientation decoder for mouse 2. The x-axis is the normalized weight of the linear SVM for discriminating img 91 (first two rows) or img 92 (last two rows); for this SVM, one orientation was held-out from img 91 and one from img 92. The weight was normalized to 1. The y-axis for odd rows is the weight of PC 1 which defines one axis of the shared plane of the relative orientation decoder; the y-axis for even rows is the weight of PC 2 which defines the other axis of the shared plane. Different columns correspond to different mouse visual cortical areas. The dashed lines highlight where weights would lie if class and orientation decoding were performed by independent neuronal populations. The decoders used here were trained on one random sample of 250 neurons in each area of mouse 2.

**Extended Data Fig 7.**
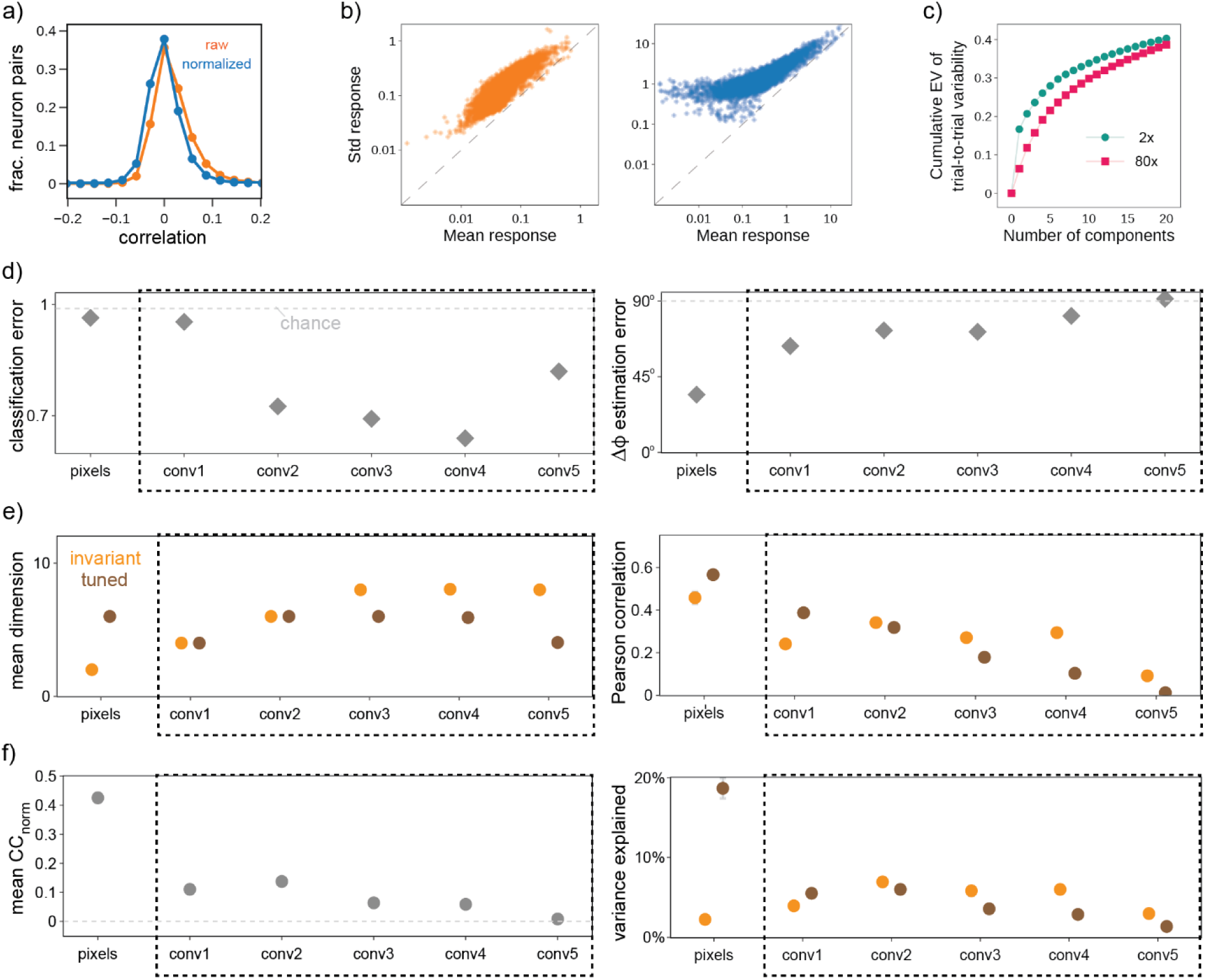
Main results for multiplicative gain model. **a)** Distribution of pairwise noise correlations between neurons before (raw, orange, mean: 0.019) and after (normalized, blue, mean: 0.002) z-scoring responses across neurons. **b)** Standard deviation versus mean response for raw (orange, left) and normalized (blue, right) data on log-log axes; dashed line indicates slope of one. c) Cumulative trial-to-trial explained variance (EV) by increasing numbers of components on 2x (teal dots) and 80x (pink squares) stimuli. Remaining panels are analogues of **d)** Fig 1d-e (*C* = 0.001), **e)** Fig 3b bottom and Fig 4b, f) Extended Data Fig 10b-c.

**Extended Data Fig 8.**
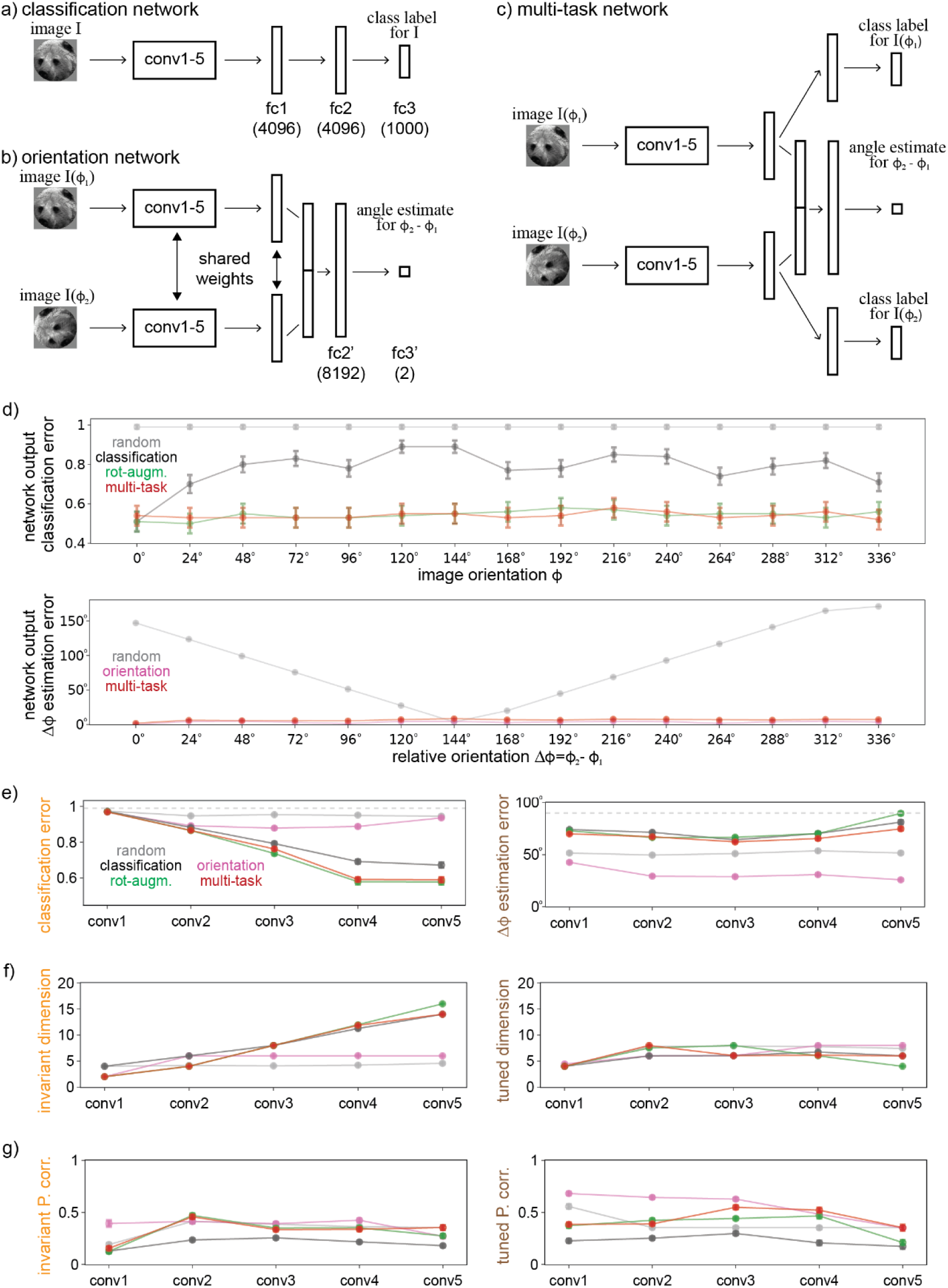
AlexNet trained on different objectives. a-c) Schematics of three variants of AlexNet. **d)** Output classification error (top) and relative orientation estimation error (bottom) as a function of orientation, for networks where the respective output is defined. Shown is mean ± sem across 100 random ImageNet test images. Remaining panels are analogues of **e)** Fig 1d-e, **f)** Fig 3b bottom, **g)** Fig 4b. The AlexNet analyses in the main text are for the “classification” network.

**Extended Data Fig 9.**
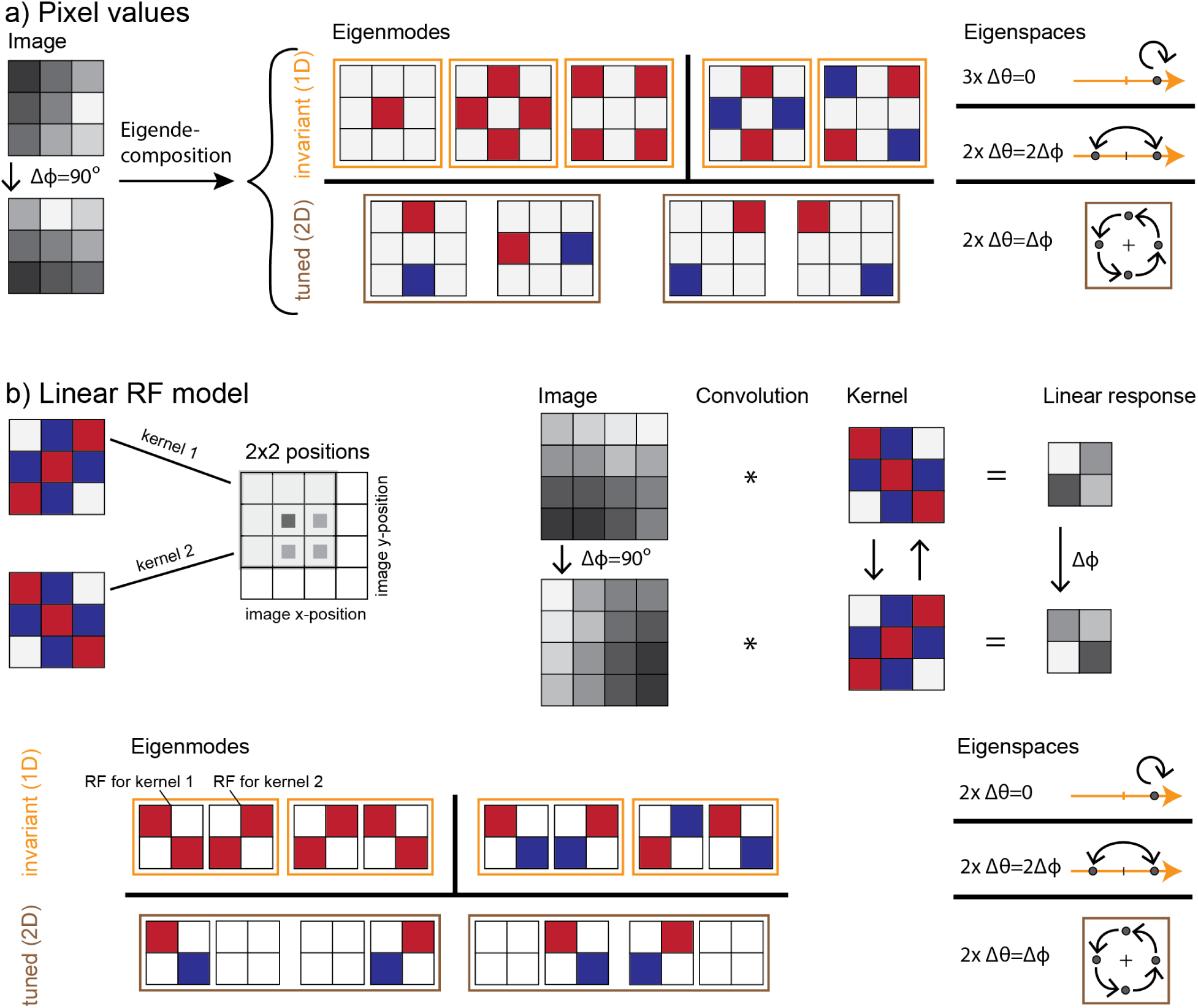
Schematic explaining rotation-equivariance for pixels and linear receptive field (RF) models, shown in. Fig 2c**. a)** Left: Example grayscale image and its Δϕ = 90° rotated version. These two images are related by a pixel permutation matrix *R*(Δϕ). Middle-right: The eigendecomposition of *R*(Δϕ) is given by its eigenmodes, the weights of which are indicated in colors blue for -1, white for 0, and red for +1, and by how they transform, which are indicated by arrows on eigenspace projections. In the eigenspaces panel, the axes are defined by the eigenmodes (in order, top-left to bottom-right), and the gray dots are the projections of the original and the rotated images to those eigenmodes. The integer before the × indicates the degeneracy, i.e., the number of eigenmodes with the same (absolute) eigenphase Δθ. **b)** Top left: The exemplified linear RF model consists of 2 kernels and 4 image positions (gray squares). The gray shaded region corresponds to the nonzero local patch of the kernel at the first image position (darker gray square). Top right: Neuronal responses are computed as convolutions of images with receptive fields (the four squares correspond to the four positions on the left). The responses of neurons with kernel 1 to the rotated image are related to the responses of neurons with kernel 2 to the unrotated image by a permutation matrix *R*(Δϕ). Bottom: The eigendecomposition of *R*(Δϕ) is analogous to that of pixel values in a, except that for the eigenmodes, one needs to specify the weights for the RFs of both kernels, shown as separate squares next to each other.

**Extended Data Fig 10.**
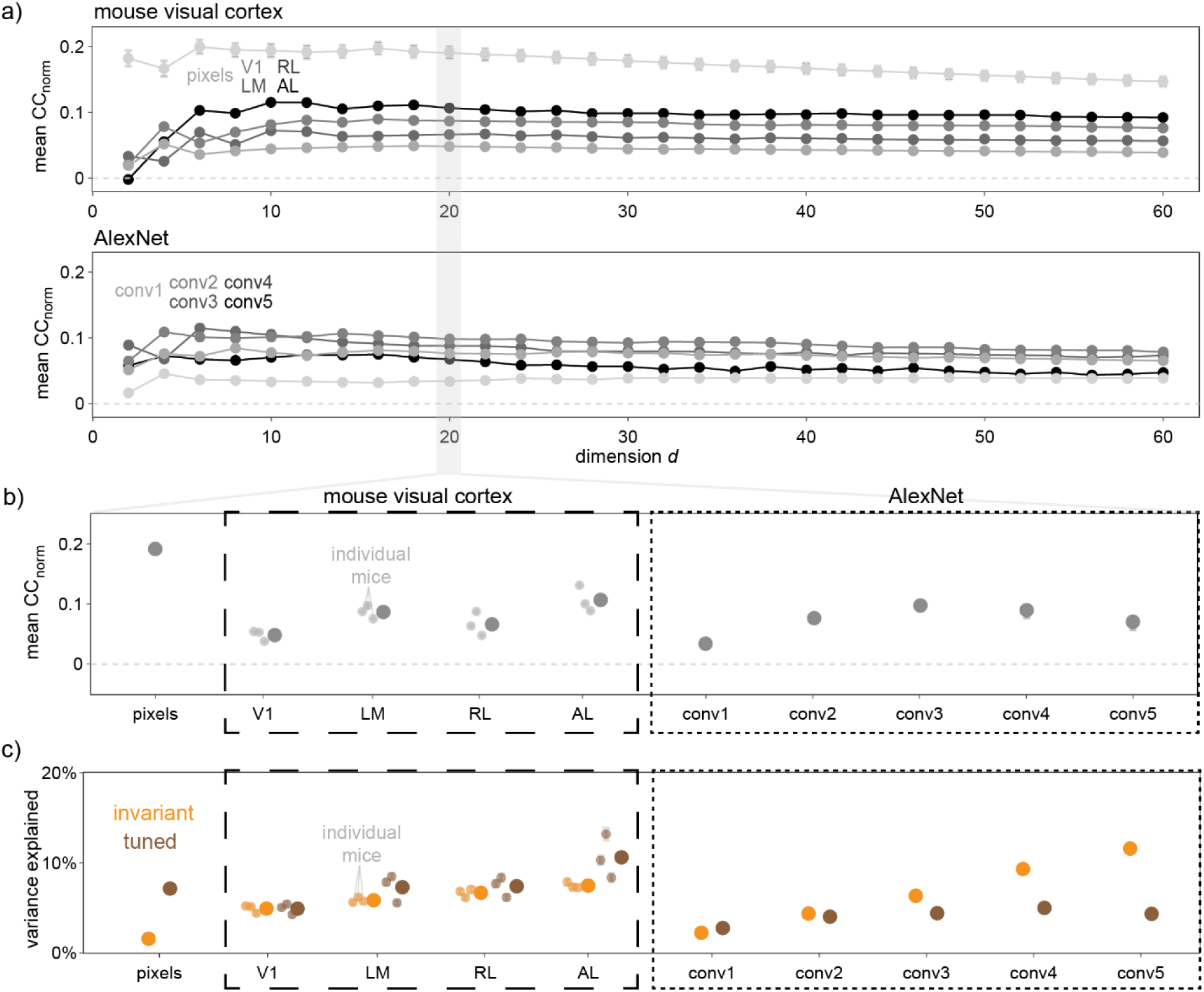
Saturation of performance of the equivariance model at dimension *d* = 20, and variance explained in the invariant and tuned subspaces. **a)** Shown is the mean normalized correlation coefficient (*CC*_*norm*_) as a function of the dimension *d* of the subspace we project neuronal responses onto. Normalization is given by the maximum possible correlation considering trial-to-trial variability. On top, different shades of gray correspond to pixels and mouse visual cortical areas, dots are means and error bars sem’s over 3 mice; on bottom, colors correspond to different AlexNet layers. Results are averaged over at most 100 independent random samples of 250 neurons. The shaded area highlights the dimension *d* = 20 we picked for most analyses. **b)** Similar to a but showing *CC*_*norm*_ only for *d* = 20. **c)** Percentages of variance explained summed over all invariant (orange) or tuned (brown) dimensions. In all panels, results were averaged over at most 100 random samples of N=250 neurons and invisible error bars are smaller than dot sizes.

