## Supplementary material for "Equivariant neuronal populations enable simultaneous tuning and invariance": SI_text

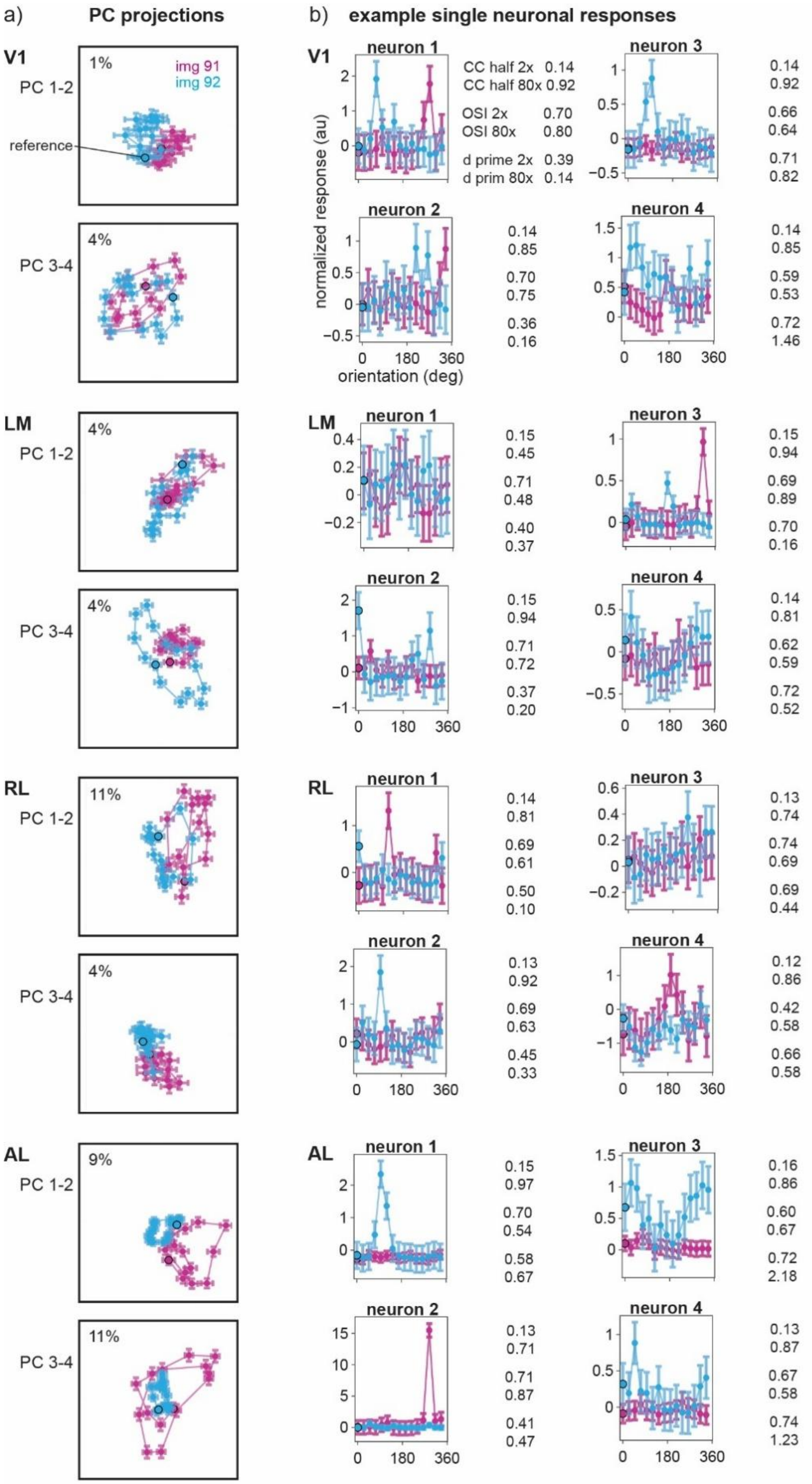


**Supplementary Fig 1. Visualization of neuronal responses across mouse visual cortical areas in mouse 1. a)** Projections of neuronal responses to the first two pairs of principal components (PCs), obtained from all reliably responsive neuronal responses to img 1–90 in each brain area (different pairs of rows). The percentage in the upper-left corner indicates the variance explained in PC 1-2 or PC 3-4 for responses to held-out images (img 91-92). Dots represent neuronal responses to img 91-92 in pink and blue, respectively. A black border indicates the reference orientation ($\phi = 0°$), and dots are connected when images differ by $\Delta\phi = 24°$. **b)** Tuning curves of example neurons in each brain area, along with values from the single-neuron distributions displayed in Extended Data Fig 1b. Example neurons were selected based on two different criteria, corresponding to the two columns. The first column shows neurons closest to the median CC half 2x and median OSI 2x (neuron 1 is the closest, neuron 2 is second-closest); the second column shows neurons closest to the median CC half 2x and median d' 2x. Colors correspond to different image classes, as in a. Dots show trial-averaged responses and error bars are sem's across 80 trials.

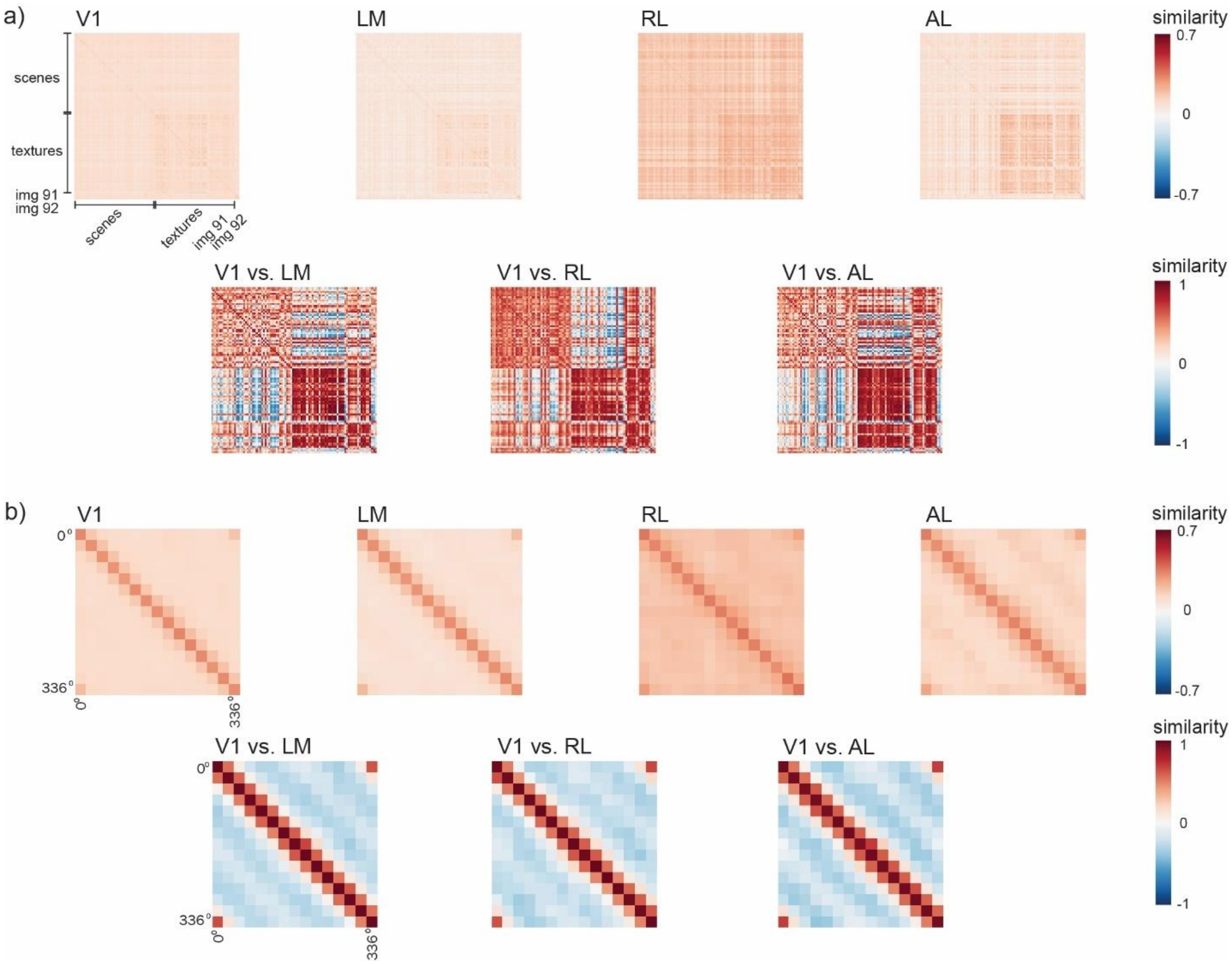


**Supplementary Fig 2. Representational similarity matrices across visual cortical areas and their correlation matrices with V1 in mouse 1. a)** Top: Representational similarity matrix for each image class, i.e., different rows/columns correspond to the different image identities. Bottom: Similarity of the matrices on top, between higher-order areas and V1. **b)** Similar to a but for each image orientation (instead of class).

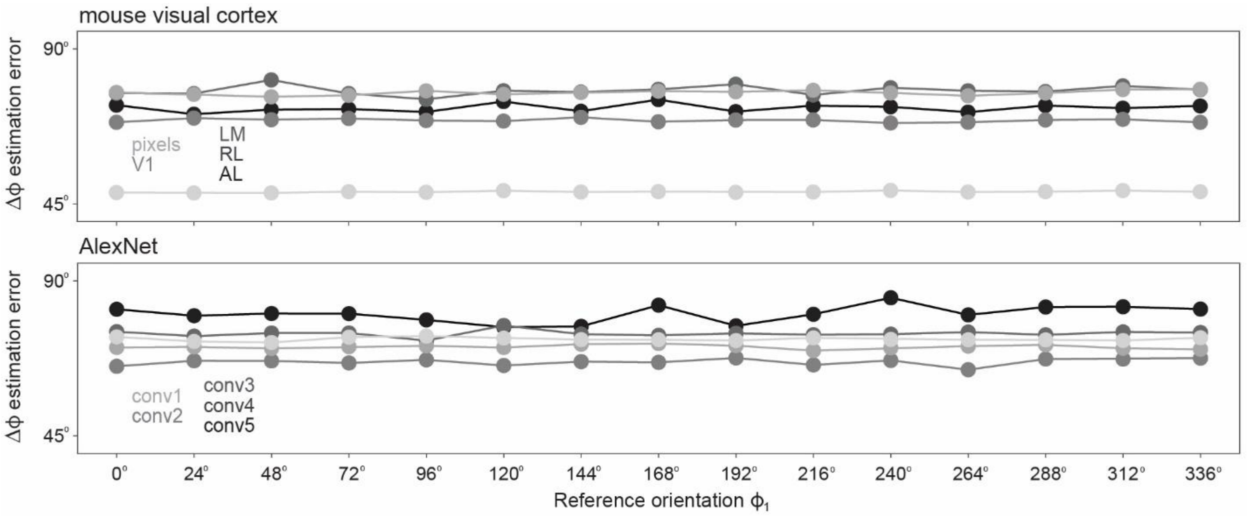


**Supplementary Fig 3. Relative orientation decoding results split by reference orientation.** Analogue of Fig 1e but for different reference orientations. Top: pixels and mouse visual cortical areas; bottom: AlexNet layers.

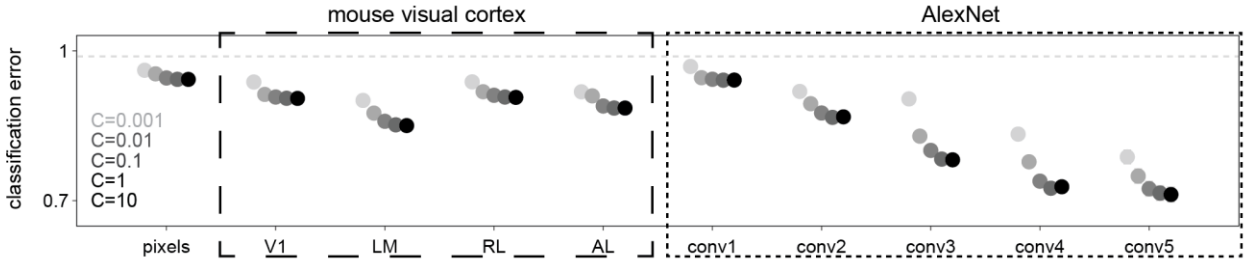


**Supplementary Fig 4. Classification results in invariant subspace for different regularization parameters C.** Analogue of Extended Data Fig 3a but for neuronal responses projected to the invariant subspace, as in Fig 3c.

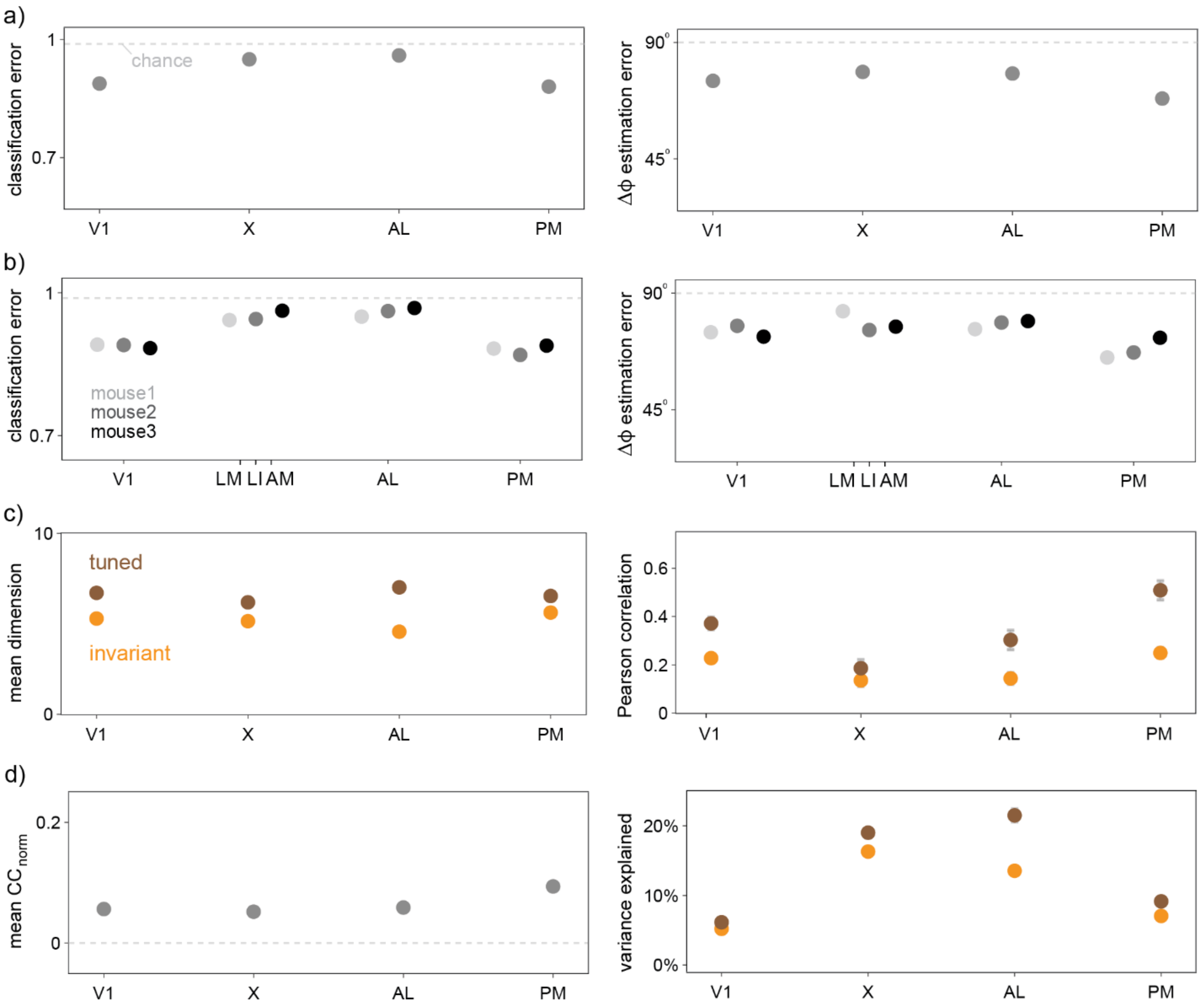


**Supplementary Fig 5. Main results for left monitor.** Analogues of **a)** Fig 1d-e, **b)** Fig 1d-e split up by mouse, **c)** Fig 3b bottom and Fig 4b, **d)** Extended Data Fig 10c for neuronal responses to images on the left-side monitor. Area X is a placeholder for areas LM, LI, AM in mouse 1, 2, 3, respectively (see Extended Data Fig 1b). Our imaging windows missed a lot of reliably responsive neurons to left-screen images in areas X and AL; hence, the results here for these areas are inconclusive. For areas V1 and PM, the imaging windows captured the majority of left-screen reliably responsive neurons, and for those areas, we used at most 100 random samples of 250 reliably responsive neurons, to enable comparison with results on the front screen.

**Supplementary data analysis methods**

*Principal component analysis*

Principal component analysis (PCA) finds the directions of largest variation of points in possibly high dimensions. We computed the PCs of trial-averaged neuronal responses to img 1-90 at all 15 orientations; this gave a total of 1350 PCs. Projections to PC 1-2 and PC 3-4 are shown in **Supplementary Fig 1a**, together with their summed variance explained. The variance explained of each PC was computed as the variance of the PC-projected neuronal responses to img 91-92 divided by the variance of the unprojected neuronal responses to 91-92. This computation did not take into account trial-to-trial variability.

*Representational similarity analysis*

Representational similarity analysis is an analysis method to quantify multivariate response patterns and compare them across sources (e.g. brain regions, behavior, computational models).[1] We first computed representational similarity matrices in each area by taking the Pearson correlation of all reliable neuronal responses at the first and at the second trial across all images (classes and orientations; for img 91-92, we selected two out of the eighty trials). For **Supplementary Fig 2a** top, we took the mean of this matrix across orientations for each class; for **Supplementary Fig 2b** top, we took the mean across classes for each orientation. To get the bottom similarity matrices, we computed the Pearson correlation of the representational similarity matrices in higher-order areas and that in V1.

*Proof that z-scoring across neurons reduces global gain*

We now show that z-scoring neuronal responses across neurons reduces global gain for small gains. The z-scored neuronal response is $z_i(t) = (r_i(t) - \bar{r}(t))/\sigma_r(t)$ where $\bar{r}(t)$ is the mean across neurons and $\sigma_r(t)$ the standard deviation. Using the expression for $r_i(t)$ in the multiplicative gain model, $r_i(t) = f_i(s_t)(1 + g_t\alpha_i)$, we find $\bar{r}(t) = \langle f_i(s_t)\rangle_i + g_t\langle f_i(s_t)\alpha_i\rangle_i$ where $\langle . \rangle_i$ denotes an average over neurons, and $\sigma_r(t)^2 = \langle f_i(s_t)^2\rangle_i + 2g_t\langle f_i(s_t)^2\alpha_i\rangle_i + g_t^2\langle f_i(s_t)^2\alpha_i^2\rangle_i - \langle f_i(s_t)\rangle_i^2 - 2g_t\langle f_i(s_t)\rangle_i\langle f_i(s_t)\alpha_i\rangle_i - g_t^2\langle f_i(s_t)\alpha_i\rangle_i^2$.

For purely global gain, $\alpha_i = \alpha$, we find that $\bar{r}(t) = \langle f_i(s_t)\rangle_i(1 + g_t\alpha)$ and $\sigma_r(t) = \sigma_f(t)(1 + g_t\alpha)$ where $\sigma_f(t)$ is the standard deviation of $f_i(s_t)$ across neurons. Therefore, $z_i(t) = (f - \langle f_i(s_t)\rangle_i)/\sigma_f(t)$ equals the z-scored $f_i(s_t)$, with zero global gain.

Let us now consider small gain, by which we mean $\alpha_i = \alpha + \delta\alpha_i$ where the global gain $\alpha$ may be large but the neuron-specific (local) factors $\delta\alpha_i$ are assumed small. Then in the numerator of $z_i(t)$, we get $r_i(t) - \bar{r}(t) = (f_i(s_t) - \langle f_i(s_t)\rangle_i)(1 + g_t\alpha) + g_t(f_i(s_t)\delta\alpha_i - \langle f_i(s_t)\delta\alpha_i\rangle_i)$, where the first term is as for the pure global gain case, and the second term is proportional to $\delta\alpha_i$ and thus small. The denominator squared, after some massaging, is $\sigma_r(t)^2 = \sigma_f(t)^2(1 + g_t\alpha)^2 + 2g_t(1 + g_t\alpha)(\langle f_i(s_t)^2\delta\alpha_i\rangle_i - \langle f_i(s_t)\rangle_i\langle f_i(s_t)\delta\alpha_i\rangle_i) + g_t^2(\langle f_i(s_t)^2\delta\alpha_i^2\rangle_i - \langle f_i(s_t)\delta\alpha_i\rangle_i^2)$, where again the first term is as for the pure global gain case, followed by a term of order $\delta\alpha_i$, and the last term will be dropped because it is of order $\delta\alpha_i^2$. Expanding to first order in $\delta\alpha_i$, we have $1/\sigma_r(t) = 1/(\sigma_f(t)(1 + g_t\alpha))(1 - g_t/(1 + g_t\alpha)\xi)$ where $\xi$ is the constant $\xi = (\langle f_i(s_t)^2\delta\alpha_i\rangle_i - \langle f_i(s_t)\rangle_i\langle f_i(s_t)\delta\alpha_i\rangle_i)/\sigma_f(t)^2$ which is proportional to $\delta\alpha_i$ and thus small. The final form of $z_i(t)$ to first order in $\delta\alpha_i$ is therefore $z_i(t) = (f_i(s_t) - \langle f_i(s_t)\rangle_i)/\sigma_f(t) + g_t/(1 + g_t\alpha)(f_i(s_t)(\delta\alpha_i - \xi) - \langle f_i(s_t)(\delta\alpha_i - \xi)\rangle_i)/\sigma_f(t)$. The first term in $z_i(t)$ is again just the z-scored $f_i(s_t)$ with zero global gain, and the second term is proportional to $g_t(\delta\alpha_i - \xi)/(1 + g_t\alpha)$, i.e., proportional to $\delta\alpha_i$, and thus small. Therefore, while $\alpha$ can lead to big changes in $r_i(t)$, the z-scored variable $z_i(t)$ is little affected by global gain $\alpha$ as long as the local gain $\delta\alpha_i$ is small.

**Rotation-equivariance**

*Definition*

Equivariance captures structure related to groups, meaning it describes transformations that have an identity, an inverse, and are associative. We first note that geometric image transformations themselves form a group; for example, planar image rotations have an identity (no rotation) and an inverse (rotation by the negative angle). If neuronal populations are equivariant, then neuronal responses change by linear transformations with similar properties as the group of image transformations itself.

Mathematically, equivariant neuronal populations are steerable,[2] or equivalently, have a linear group representation of geometric image transformations.[3] Let us formalize what this means. We may think of visual neuronal population (en)coding as mapping images into neuronal population responses, i.e., constituting a map $f: \mathfrak{I} \to \mathfrak{R}$ where $\mathfrak{I}$ is the image space and $\mathfrak{R}$ the neuronal response space. Images taken by a perspective camera may be modeled as the projective plane, $\mathbb{R}P^2$, and images of a planar surface taken from different 3D viewpoints are linearly related to each other by 2D projective transformations, $\mathrm{PGL}_3(\mathbb{R})$.[4] These transformations are the geometric image transformations, and $\mathfrak{I}$ has a linear group representation of $\mathrm{PGL}_3(\mathbb{R})$. If $I_1$ is the image of a planar surface taken from one viewpoint, then an image $I_2$ taken from a different viewpoint can be simply obtained from $I_1$ via a matrix-vector multiplication, which we denote as $I_2 = g \cdot I_1$, where the group element $g$ of $\mathrm{PGL}_3(\mathbb{R})$ encodes the change in viewpoint.

For a visual neuronal population to be equivariant formally means that there exists a linear group representation $R$ on $\mathfrak{R}$ such that $f(g \cdot I) = R(g) \cdot f(I)$. This means that there exists another map $R: \mathrm{PGL}_3(\mathbb{R}) \to GL(\mathfrak{R})$ such that $R(g_1 g_2) = R(g_1)R(g_2)$, i.e., $R$ is a group homomorphism ("compositional"). To clarify, $R(g)$ is a matrix acting in neuronal response space, describing how neuronal responses change as the viewpoint changes via $g$. In the mathematics literature, equivariance is typically studied in the context of linear maps which are fully characterized by Schur's lemma. But for visual neuronal responses, $f$ is typically nonlinear. A sufficient condition for equivariance is that $f$ is invertible.[5]

*Orthogonality of $R(\Delta\phi)$*

In the main text we considered in-plane rotations only, i.e., $R: \mathrm{SO}(2) \to GL(\mathfrak{R})$ where an element $g$ the group of 2D rotations $\mathrm{SO}(2)$ is parametrized by an angle $\phi$, i.e., $g \equiv g_\phi$, and we shall use the notation $R(\phi)$ instead of $R(g_\phi)$. Let $r(I, \phi) = f(g_\phi \cdot I) \in \mathfrak{R}$ be the neuronal population response to image $I$ at orientation $\phi$. We now show that if orientations $\phi$ are sampled uniformly, as in the main text, then after whitening the neuronal responses, $R$ is orthogonal. The whitening transform is defined as setting covariance matrix to the identity; it is used, for instance, in canonical correlation analysis.[6]

We first note that the law of total covariance shows that the covariance of neuronal responses can be decomposed into within-image and between-image covariances, i.e., $\Sigma = \mathrm{cov}_{I,\phi}\big(r(I,\phi), r(I,\phi)\big) = \mathbb{E}_I\big[\mathrm{cov}_\phi\big(r(I,\phi), r(I,\phi)\big)\big] + \mathrm{cov}_I\big(\mathbb{E}_\phi[r(I,\phi)], \mathbb{E}_\phi[r(I,\phi)]\big)$. Each term is SO(2)-invariant: the within-image covariance because shifting $\phi$ to $\phi + \phi'$ by any $\phi'$ in the sum leaves it unchanged under uniform sampling, and the between-image covariance because the image-mean is itself invariant under shifting. Therefore, the total covariance $\Sigma$ is also SO(2)-invariant, which means that $R(\phi)\Sigma R(\phi)^T = \Sigma$ and

$R(\phi)^T\Sigma^{-1}R(\phi) = \Sigma^{-1}$ (by inverting both sides and replacing $\phi$ by $-\phi$) for all $\phi$. This construction is dual to what is known as the unitary trick.

Next, we consider the whitening transformation $\tilde{r} = \Sigma^{-1/2}r$. The matrix in the whitened basis becomes $\tilde{R}(\phi) = \Sigma^{-1/2}R(\phi)\Sigma^{1/2}$ which satisfies $\tilde{R}(\phi)^T\tilde{R}(\phi) = \Sigma^{1/2}(R(\phi)^T\Sigma^{-1/2}\Sigma^{-1/2}R(\phi))\Sigma^{1/2} = \Sigma^{1/2}\Sigma^{-1}\Sigma^{1/2}$ where we used SO(2)-invariance of $\Sigma^{-1}$. This shows that $\tilde{R}(\phi)^T\tilde{R}(\phi)$ equals the identity matrix (and similarly for the transpose) and thus $\tilde{R}(\phi)$ is orthogonal.

In our preprocessing of neuronal responses, we z-score across neurons (reducing shared multiplicative-gain correlations across neurons) and divide each neuronal response by its standard deviation ("diagonal whitening", i.e., rescaling each neuron to unit variance). Full whitening is sample-inefficient and often approximated by diagonal whitening alone in neuroscience; our two preprocessing steps together provide a closer approximation of full whitening. The orthogonality of $\tilde{R}(\phi)$ is enforced by the Procrustes constraint and is therefore independent of whether the preprocessed neuronal responses were exactly whitened. The residual fit error reflects a combination of imperfect equivariance, imperfect whitening, and noise; it is quantified by the Pearson correlations between the data and the equivariance model in the invariant and tuned subspaces (**Fig 4b**).

*Eigendecomposition of orthogonal $R(\Delta\phi)$*

A (real) eigendecomposition of an orthogonal $m \times m$ matrix $R(\Delta\phi)$ decomposes $m$-dimensional neuronal response space into mutually orthogonal one- and two-dimensional eigenspaces, which are spanned by the eigenmodes (or eigenvectors) of $R(\Delta\phi)$. Formally, there exists an orthogonal $m \times m$ matrix $Q$ (the columns of which are the eigenmodes) such that

$$R(\Delta\phi) = Q \begin{pmatrix} R_{2D}(\Delta\theta_1) & & & & & \\ & \ddots & & & & \\ & & R_{2D}(\Delta\theta_k) & & & \\ & & & \pm 1 & & \\ & & & & \ddots & \\ & & & & & \pm 1 \end{pmatrix} Q^T$$

where $R_{2D}(\Delta\theta_j) = \begin{pmatrix} \cos(\Delta\theta_j) & -\sin(\Delta\theta_j) \\ \sin(\Delta\theta_j) & \cos(\Delta\theta_j) \end{pmatrix}$ is a simple 2D rotation by an angle $\Delta\theta_j$, $j = 1, \ldots, k$ for $0 \leq k \leq \lfloor m/2 \rfloor$, and all empty entries are 0. This means that $R(\Delta\phi)$ either maps a single eigenmode onto itself (times $\pm 1$) or a pair of eigenmodes onto themselves via a 2D rotation $R_{2D}(\Delta\theta_j)$—such pairs of eigenmodes are also known as quadrature pairs.[7] The angles $\Delta\theta_j$ are the eigenphases of $R(\Delta\phi)$, so-called because the eigenvalues of $R(\Delta\phi)$ have the form $e^{\pm i\Delta\theta_j}$ (a direct consequence of orthogonality).

*Additional constraints on $R(\Delta\phi)$*

Compositionality and periodicity constraints on $R(\Delta\phi)$ imply that the $\Delta\theta_j$ are integer multiples of the image rotation angle $\Delta\phi$. This can be seen by setting $\Delta\phi = 2\pi/n$ for an integer $n > 1$, noting that compositionality implies that $R(\Delta\phi)^n = R(n\Delta\phi) = R(2\pi)$ and periodicity implies that $R(2\pi) = R(0)$ equals the identity matrix. Hence, the eigenvalues $e^{\pm i\Delta\theta_j}$ must satisfy $e^{\pm in\Delta\theta_j} = 1$, which means that $n\Delta\theta_j$ must be a multiple of $2\pi$, i.e., $\Delta\theta_j$ is a multiple of $2\pi/n = \Delta\phi$.

*Numerically estimated eigenspaces*

In numerical computations, one-dimensional eigenspaces are rare because that would require estimating the eigenvalues $\pm 1$ exactly. In odd dimensions, one such one-dimensional eigenspace has to exist. In our numerical computations, we found this typically to be the only one-dimensional eigenspace. To avoid the unnecessary complexity in having one one-dimensional eigenspace, we only considered even dimensions in the main text, where all our eigenspaces were two-dimensional. For eigenspaces with $\Delta\theta \approx 0°$ and $\Delta\theta \approx 180°$, the transformation induced by $R(\Delta\phi)$ does not mix the corresponding pair of eigenmodes much and either maps points onto themselves or onto minus themselves; therefore, we called such eigenspaces "invariant". For $\Delta\theta \approx \pm\Delta\phi$, the transformation mixes the corresponding pair of eigenmodes and acts on points like a planar rotation by $\Delta\phi$. These constitute the "tuned" eigenspaces.

There exist eigenspaces for which the corresponding eigenphases are higher-order multiples of $\Delta\phi$, which we did not discuss further as they were not relevant for our simple decoders of image identity and relative orientation.
